# Multi-dimensional profiling of hepatoblastomas and patient-derived tumor organoids uncovers tumor subpopulations with divergent WNT activation profiles and identifies pan-hepatoblastoma drug sensitivities

**DOI:** 10.1101/2023.08.28.554783

**Authors:** Thomas A. Kluiver, Yuyan Lu, Stephanie A. Schubert, Lianne J. Kraaier, Femke Ringnalda, Philip Lijnzaad, Jeff DeMartino, Wout L. Megchelenbrink, Vicky Amo-Addae, Selma Eising, Flavia W. de Faria, Daniel J. Münter, Marc van de Wetering, Kornelius Kerl, Evelien Duiker, Marius van den Heuvel, Vincent E. de Meijer, Ruben H. de Kleine, Ronald R. de Krijger, Jan J. Molenaar, Thanasis Margaritis, Henk Stunnenberg, József Zsiros, Hans Clevers, Weng Chuan Peng

**Affiliations:** Princess Máxima Center for Pediatric Oncology, Heidelberglaan 25, 3584 CS, Utrecht, the Netherlands; Department of Hepatobiliary Surgery, Xiamen Hospital of Traditional Chinese Medicine, Beijing University of Chinese Medicine, Xiamen, China; Department of Pediatric Hematology and Oncology, University Children’s Hospital Münster, Albert-Schweitzer-Campus 1, 48149, Münster, Germany; Department of Surgery, Section of Hepatobiliary Surgery and Liver Transplantation, University of Groningen, University Medical Center Groningen, Groningen, the Netherlands; Department of Pathology, University Medical Center Utrecht, Heidelberglaan 100, 3584 CX, Utrecht, the Netherlands; Hubrecht Institute, Royal Netherlands Academy of Arts and Sciences and University Medical Center, 3584 CT, Utrecht, the Netherlands; Oncode Institute, the Netherlands

## Abstract

Hepatoblastoma, the most prevalent pediatric liver cancer, almost always carries a WNT-activating *CTNNB1* mutation, yet exhibits notable molecular heterogeneity. To characterize this heterogeneity and identify novel targeted therapies, we performed comprehensive analysis of hepatoblastomas and tumor-derived organoids using single-cell RNA-seq, spatial transcriptomics, single-cell ATAC-seq and high throughput drug profiling. We identified two distinct tumor epithelial signatures: hepatic ‘fetal-like’ and WNT-high ‘embryonal-like’ signatures, displaying divergent WNT signaling patterns. The liver-specific WNT targets were enriched in the fetal-like group, while the embryonal-like group was enriched in canonical WNT target genes. Gene regulatory network analysis revealed enrichment of regulons related to hepatic function such as bile acid, lipid and xenobiotic metabolism in the fetal-like subgroup but not in the embryonal-like subgroup. In addition, the dichotomous expression pattern of the transcription factors HNF4A and LEF1 allowed for a clear distinction between the fetal- and embryonal-like tumors. We also performed high-throughput drug screening using patient-derived tumor organoids and identified sensitivity to multiple inhibitor classes, most notably HDAC inhibitors. Intriguingly, embryonal-like tumor organoids, but not fetal-like tumor organoids, were sensitive to FGFR inhibitor treatments, suggesting a dependency on FGFR signaling. In summary, our data uncover the molecular and drug sensitivity landscapes of hepatoblastoma and pave the way for the development of targeted therapies.

## INTRODUCTION

Hepatoblastoma, the most prevalent liver malignancy in children, is slowly increasing in incidence^1–3^. Although cisplatin-based chemotherapy regimens and surgical resection have improved survival^4–6^ for patients with low-risk tumors, the prognosis of children with advanced or high-risk hepatoblastoma (60% of all patients) remains unsatisfactory due to a high recurrence rate and progressive disease due to the development of chemoresistance^7^. Moreover, long-term side effects from chemotherapy, such as hearing loss, reduced cardiac and renal function, and secondary cancers, can severely impact childhood development and quality of life in survivors^8^. Improvements in care for these patients require the development of targeted therapeutic treatments.

Hepatoblastomas have a low mutational burden and few recurrent chromosomal aberrations^9,10^. Notably, they are characterized by the presence of activating mutations in *CTNNB1*, which encodes β-catenin, or in other WNT pathway genes, accounting for over 90% of cases^2^. These tumors are believed to originate from aberrant expansion of hepatic progenitors that harbor *CTNNB1* mutations during fetal liver development, and morphologically resemble stages of liver development^11^. Within the tumor epithelium, two main histological types are commonly observed: the more differentiated ‘fetal’ histology, which resembles the fetal liver developmental stage, and the less differentiated ‘embryonal’ histology, resembling the earlier stage of liver development. Often, tumors can contain both fetal and embryonal histological components. It is generally hypothesized that tumors enriched in embryonal components are associated with worse prognosis^12^.

The molecular profile of hepatoblastoma has been characterized previously by several groups using bulk methods^13–19^. These studies, based on different cohorts, generally agree with each other on the existence of three main molecular subgroups within hepatoblastoma, *i.e.*, tumors with a predominantly differentiated fetal histology, enriched in a hepatic signature, tumors with a predominant embryonal histology, enriched in a progenitor signature, and a third group characterized by a mesenchymal signature^14,16–21^. More recently, single-cell transcriptomics studies based on a small cohort of patients reported a classification in five^22^ or seven tumor cell clusters^19^. Additionally, several studies have reported biological markers that showed a correlation with clinical behavior and outcome, such as the 16-gene signature, four protein markers, expression of vimentin, and the 14q32-gene signature^14,15,18,20,21,23^. However, the transcriptomic heterogeneity observed in hepatoblastoma, especially at the single-cell level, remains largely unexplored.

In this study, we investigated hepatoblastomas and organoids derived from patient tumor material using single-cell RNA-seq (scRNA-seq), spatial transcriptomics (ST) and single-cell assay for transposase-accessible chromatin (scATAC-seq) techniques. Focusing on the tumor epithelial component, we identified two distinct tumor subpopulations, which we denoted ‘fetal-like’ and ‘embryonal-like’. Fetal-like tumor cells expressed hepatic markers, including fetal liver and pericentral hepatic markers, while embryonal-like cells are enriched in WNT pathway-related marker expression. Patient-derived organoids recapitulated tumor heterogeneity, facilitating high-throughput drug screening, and revealing sensitivities to various inhibitor classes, such as HDAC inhibitors, proteasome inhibitors, PLK-1 inhibitors and FGFR inhibitors. Interestingly, fetal- and embryonal-like tumor organoids displayed different sensitivity profiles to some inhibitors, with FGFR inhibitors specifically targeting embryonal-like tumor organoids. To our knowledge, this study represents one of the first extensive screening efforts using a large cohort of patient-derived organoids of pre- and post-treatment tumors, providing essential insights, and opening new avenues for targeted therapy for children with liver tumors.

## RESULTS

### scRNA-seq confirmed the presence of fetal- and embryonal-like signatures in hepatoblastoma

To investigate the heterogeneity of hepatoblastoma tumor cells, we analyzed a recently published scRNA-seq dataset of nine hepatoblastomas from Song *et al.*^22^ (**Figure S1A, B)**. We focused on the epithelial tumor cells, while the mesenchymal component and the very rare tumor cluster with neuroendocrine features were excluded from our analysis. In addition, we included hepatocytes and cholangiocytes from the paired normal tissue for comparison. Single-cell data was processed as described in Experimental Procedures. In total, 1562 cells from 9 patients were jointly analyzed. Based on unsupervised graph-based clustering, normal hepatocytes and cholangiocytes from multiple patients were present as two separate clusters **(Figure 1A)**. From tumor tissues, we identified two distinct clusters of cells originating from multiple patients.

**Figure 1.**
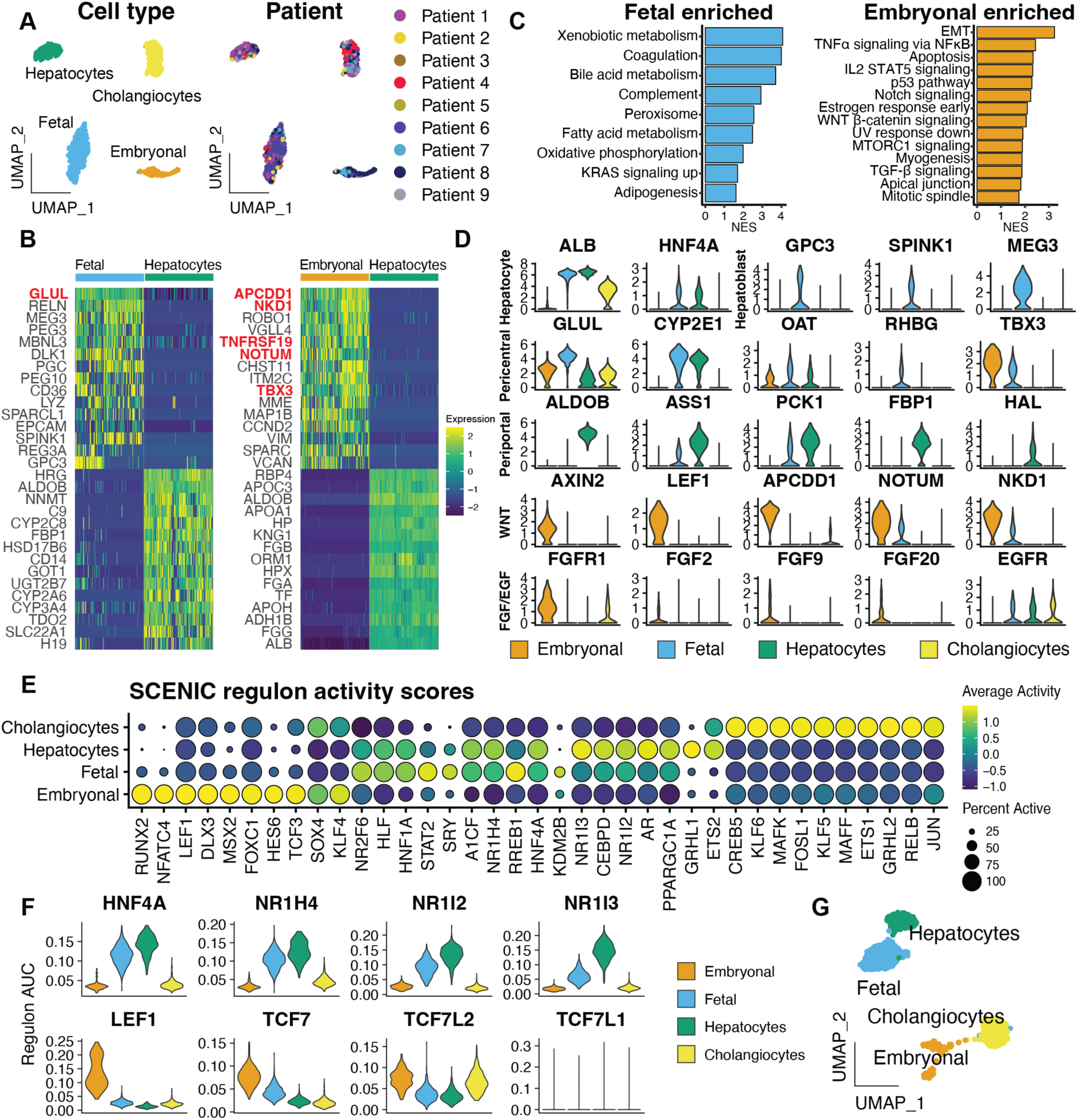
ScRNA-seq analysis of primary tumor material illustrates fetal- and embryonal-like tumor signatures. (A) UMAP plots based on unsupervised clustering, annotated per cell type or tumor signature (left) and patient identity (right) of epithelial cell subsets from the Song *et al.* dataset. (B) Heatmaps showing the top differentially expressed genes across tumor cell populations and hepatocytes. WNT signaling pathway target genes are upregulated especially in the WNT-high embryonal-like tumor subpopulation and marked in red. (C) Gene set enrichment analysis showing hallmark gene sets enriched in the fetal- and embryonal-like clusters, compared to each other. NES: normalized enrichment score. (D) Violin plots showing expression of select hepatic and WNT-related markers. (E) Scaled dot plot showing the most differentially enriched transcription factor regulons per cell type. (F) Violin plots showing regulon activity scores for hepatic TFs (above) enriched in fetal-like tumor cells and WNT/β-catenin transcription factors (below) in embryonal-like tumor cells. (G) UMAP plot based on the SCENIC regulon activity scores.

We first examined the tumor expression patterns compared to normal hepatocytes and discovered that one tumor cluster exhibited high levels of hepatocyte specific WNT target genes, *i.e.*, pericentral markers (*e.g., GLUL, CYP2E1* and *RHBG)*^24^ with low levels or absence of various periportal hepatic markers (*e.g., ALDOB*, *PCK1* and *FBP1*) **(Figure 1B**, D**, S1C)**. These cells also expressed other general WNT target genes, such as *NOTUM* and *NKD1*, but at a lower level than the other tumor cluster. Additionally, these tumor cells upregulated fetal liver markers (*e.g., SPINK1*, *GPC3*, *REG3A* and *RELN*)^14,25–28^, and expressed multiple imprinted genes, including *IGF2*, *PEG3*, *PEG10, DLK1* and *MEG3* **(Figure 1B, S1C)**, with *DLK1* and *MEG3* being located on the 14q32 locus, previously described by Carrillo-Reixach *et al.*^21^. This expression profile is consistent with previously described tumors enriched with a hepatic signature^14,16–19^, which we denote here ‘fetal-like’ (F). These tumors cells were described as Hepatic I and II in Song *et al.*^22^.

We then compared the second tumor cluster to normal hepatocytes and observed significant upregulation of key WNT pathway target genes including *APCDD1*^29^*, NKD1*, *NOTUM, KREMEN1, TNFRSF19* and *AXIN2*, as well as WNT target genes that are associated with epithelial-to-mesenchymal transition (EMT) such as *TWIST1, BMP4* and *VIM*^30^, along with low levels of hepatic markers **(Figure 1B-D, S1C)**. This WNT-high expression profile is consistent with previously described tumors with a progenitor signature^16–21^, which we denote here ‘embryonal-like’ (E). These tumor features were not described in the original study^22^.

Next, using gene set enrichment analysis, we compared the fetal- and embryonal-like tumor cells with each other **(Figure 1C)**. Fetal-like tumor cells were enriched for hepatic functions, such as xenobiotic and bile acid metabolism, coagulation, and the complement system. In turn, the embryonal-like cells were enriched for EMT, WNT/β-catenin signaling, the P53 pathway, and mitotic spindle-related genes **(Figure 1C)**. The embryonal-like tumor cluster showed significantly lower expression levels of hepatic markers relative to both fetal-like tumor cells and normal hepatocytes **(Figure 1D)**. Interestingly, we identified differential expression of FGFR1 and several FGF ligands in the embryonal-like tumor cluster, and EGFR in the fetal-like tumor cluster, suggesting a potential role of the EGF/FGF signaling pathways in hepatoblastoma tumorigenesis **(Figure 1D)**. In brief, consistent with previous bulk RNA-seq studies^13–19^, our scRNA-seq analysis revealed the presence of two distinct profiles in hepatoblastoma cells: a fetal-like profile with hepatic features, pericentral and fetal liver markers, and an embryonal-like profile with high expression of WNT pathway genes and reduced hepatic markers^12–18^.

### Fetal- and embryonal-like tumor cells are enriched in hepatic and WNT pathway-related regulons, respectively

To uncover the gene regulatory network underlying the different hepatoblastoma subpopulations, we employed single-cell regulatory network inference and clustering (SCENIC)^31^ analysis. SCENIC utilizes scRNA-seq co-expression patterns in combination with *cis*-regulatory motif analysis to infer the activity of transcription factors (TFs) and their target genes (termed ‘regulon activity’). Gene regulatory network analysis has the potential to better distinguish cellular heterogeneity than RNA-seq alone, as it identifies key transcriptional activity (TFs) underlying a specific cellular state^31^.

Consistent with previous studies^32^, our analysis revealed that cholangiocytes were enriched for regulons such as ONECUT1, HNF1B and SOX9 **(Figure 1E)**. The fetal-like tumor cluster displayed enrichment for hepatic-specific regulons. These included hepatic nuclear factor 4A (HNF4A), FOXA3 (*i.e.*, HNF3G), androgen receptor (AR), and multiple nuclear receptor subfamily members such as NR1H4 (*i.e.,* farnesoid X-activated receptor [FXR]), NR1I3 (*i.e.,* constitutive androstane receptor [CAR]) and NR1I2 (*i.e.,* pregnane X receptor [PXR])) **(Figure 1E, 1F)**. Collectively, these TFs regulate essential hepatic functions such as bile acid, lipid, and xenobiotic metabolism, and were also present in normal hepatocytes **(Figure 1E, 1F, S1D).** Notably, some of these TFs, such as CAR and PXR, are activated by the WNT pathway^33,34^.

In contrast, the embryonal-like tumor cluster mostly lacked these hepatic TFs, or they were expressed at minimal levels. This might explain the observe low levels of hepatic markers in this subgroup (**Figure 1C, 1D**). However, this cluster showed a clear enrichment of WNT pathway regulons^35–38^, such as LEF1, TCF7, TCF7L2 and MSX2 **(Figure 1E, 1F**). This correlates with the observed high expression of WNT target genes (**Figure 1C, 1D**). Although the fetal-like tumor cluster displayed some regulon activity for TCF7/TCF7L2, it was notably less pronounced compared to the embryonal-like tumor cluster.

Next, we performed unsupervised graph-based clustering using the inferred regulon activity scores and visualized this in a UMAP plot **(Figure 1G)**. Hepatocytes and cholangiocytes, irrespective of patient origin, formed two distinct clusters, serving as a reference for comparison. Similarly, the fetal-like tumor cells from different patients clustered closely together, suggesting a high degree of similarity. In contrast, the embryonal-like cells formed separate clusters that were associated with specific patient origin, suggesting the presence of inter-patient heterogeneity **(Figure S1D)**. Overall, our analysis uncovered distinct gene regulatory network underlying fetal- and embryonal-like tumor clusters, highlighting the roles of hepatic-specific and WNT pathway related TFs in driving these distinct tumor features observed in RNA-seq.

### Spatial molecular landscape of hepatoblastoma reveals tumor heterogeneity

To study the *in-situ* expression patterns of hepatoblastoma, we performed ST analysis using the *10x Genomics Visium* platform on four hepatoblastoma tissues, as well as one paired normal liver tissue sample (**Figure 2A**). The clinical information is provided in **Table S1**. Using this platform, each ST spot, measured at 55 μm, contained multiple cells (5-50 cells). ST spots were clustered using unsupervised graph-based clustering and annotated into different regions based on marker gene expression, differential gene expression and tissue histology.

**Figure 2.**
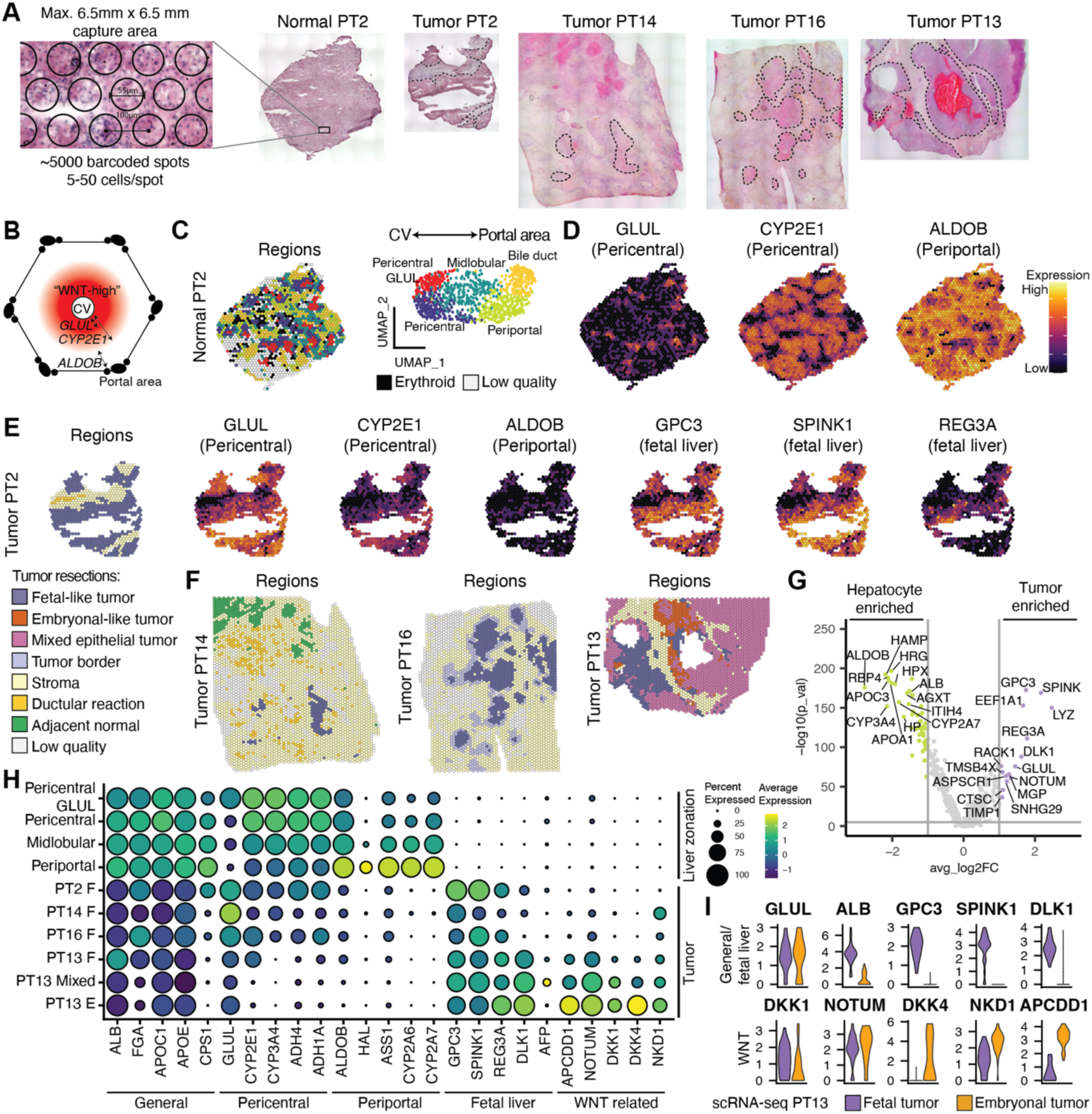
Spatial transcriptomics of hepatoblastoma tissues displays distinct spatial molecular patterns. (A) ST analysis was performed using the *10x Genomics Visium* platform on four hepatoblastoma tissues, as well as one paired distal normal liver tissue. H&E staining tissue sections are shown, with the tumor regions indicated (dotted line). Schematic representation of a liver lobule showing a “WNT-high” gradient around the central vein (CV), associated with high (B) *GLUL* (narrow) and *CYP2E1* (broad) expression, while the periportal zone is associated with a distinct hepatic expression profile, including expression of genes such as *ALDOB*. (C) Spatial map (left) and UMAP (right) annotated by liver metabolic zonation clusters. (D) Spatial map of liver zonation markers showing distinct expression patterns of pericentral and periportal markers in normal liver. (E) Spatial map of clusters and liver markers showing retain (reduced) expression of pericentral markers, reduced expression of periportal markers and expression of fetal liver markers in tumor regions of PT2, but absent in stromal regions. (F) Spatial map of clusters in three additional hepatoblastoma resections (PT14, PT16, PT13), showing fetal-like tumor regions. Additional heterogeneity is observed in PT13, where we identified fetal-like, embryonal-like and mixed tumor regions. (G) Volcano plot of differentially expressed genes between all tumor regions and distal hepatocytes (pericentral, midlobular and periportal) highlighting the expression of tumor-specific fetal liver markers and reduced expression of mature hepatic and periportal markers. (H) Dot plot comparing expression of hepatic markers between hepatocyte regions and tumor regions. (I) Violin plot showing two distinct tumor populations in scRNA-seq data of PT13 tumor resection, corresponding to the fetal- and embryonal-like tumor spots observed in the ST analysis.

Within normal liver, liver metabolic zonation could readily be distinguished, with *GLUL* expression in hepatocytes adjacent to endothelial cells of the central veins and periportal markers, such as *ALDOB*, expressed in hepatocytes adjacent to the portal triad^39,40^ (**Figure 2B**). As expected, in the section of distal normal liver of PT2, we could distinguish zonation patterns, including pericentral, midlobular, periportal and bile duct regions (**Figure 2C, S2A-C**). We observed regions with restricted *GLUL* expression and broader *CYP2E1* expression, annotated as ‘GLUL pericentral’ and ‘pericentral’, respectively (**Figure 2D**). In the periportal region, we observed expression of periportal markers *(e.g., ALDOB*, *HAL*, and *CYP2A7*) (**Figure 2D, S2C, S2D**).

We analyzed the matched tumor tissue of PT2 **(Figure 2E, S2E-G**), mostly consisted of fetal hepatoblastoma (annotated as ‘fetal-like tumor’) and stromal regions (containing ‘stroma’ and ‘ductular reaction’), which were confirmed by a pathologist (RRdK). We compared the gene expression profile of the tumor cluster with the non-tumor clusters and paired distal normal tissue section (**Figure S2H)**. In line with our scRNA-seq tumor signatures, the fetal-like tumor region upregulated fetal liver and tumor markers (*e.g.*, *GPC3*, *SPINK1*, *REG3A*), which were not detected in the normal liver (**Figure 2E, S2D, S2E**). In addition, pericentral hepatic markers and WNT target genes, such as *CYP2E1* and *GLUL*, were broadly expressed in the tumor region, contrasting with the zonated pattern in the normal tissue (**Figure 2D, 2E**). Conversely, hepatic periportal markers (*e.g.*, *ALDOB*, *HAL*, *ASS1*) and a subset of *CYP* related proteins (*e.g.*, *CYP2A6*, *CYP2A7*, *CYP2B6*) were either absent or downregulated in the tumor region (**Figure S2H**).

We analyzed three additional tumor sections (PT13, PT14 and PT16; **Figure S3A-C**). All samples, except PT13, were collected post-chemotherapy. In these sections, we identified tumor, stromal and normal hepatocyte regions (**Figure 2F**). Consistent with PT2, tumor regions from PT13, PT14 and PT16 showed similar tumor-specific gene expression profiles compared to hepatocytes from the distal normal tissue, with broad expression of pericentral hepatic and fetal markers and lower level of periportal markers (**Figure 2G, 2H, S3D**). Of note, PT13 showed considerably more heterogeneity compared to the remaining post-chemotherapy tumors. Some tumor regions displayed high levels of WNT target genes (*e.g.*, *NOTUM*, *DKK4*, *NKD1, APCDD1*) and lower levels of hepatic markers (**Figure 2H, S3A**), in line with the embryonal-like signature and was therefore annotated as such. However, the bulk of the tumor showed expression of both fetal- and embryonal-like tumor profiles, indicating ST spots with a mixture of the two epithelial tumor cell types. We confirmed the presence of both tumor populations using scRNA-seq (**Figure 2I**). Across the four tumor samples, inter-tumoral heterogeneity was also observed (**Figure S3E**). Overall, our ST analysis is consistent with fetal- and embryonal-like tumor signatures, and further reveals heterogeneity in the spatial context.

### HNF4A and LEF1 marked distinct tumor subpopulations

To validate the presence of fetal- and embryonal-like tumor clusters identified through our scRNA-seq and spatial analyses, we conducted immunofluorescence staining on a limited series of FFPE tumor samples collected from 9 patients **(Table S1)**. Based on the results of our gene regulatory network analysis, we first stained for the TFs HNF4A, the central hepatic regulator in the liver^41^, and LEF1, a key mediator in the WNT/β-catenin pathway^35^. We were able to identify tumor cells which stained strongly for either HNF4A or LEF1, with the staining pattern of the two TFs being mutually exclusive **(Figure 3)**. We observed separate regions of HNF4A^+^ cells and LEF1^+^ tumor cells, but also LEF1^+^ cells that were interspersed with HNF4A^+^ cells, as illustrated in PT13. Of note, islands of LEF1^+^ cells separated by stromal cells from surrounding HNF4A^+^ cells were also observed in post-chemotherapy samples (PT9, 17) (**Figure 3**).

**Figure 3.**
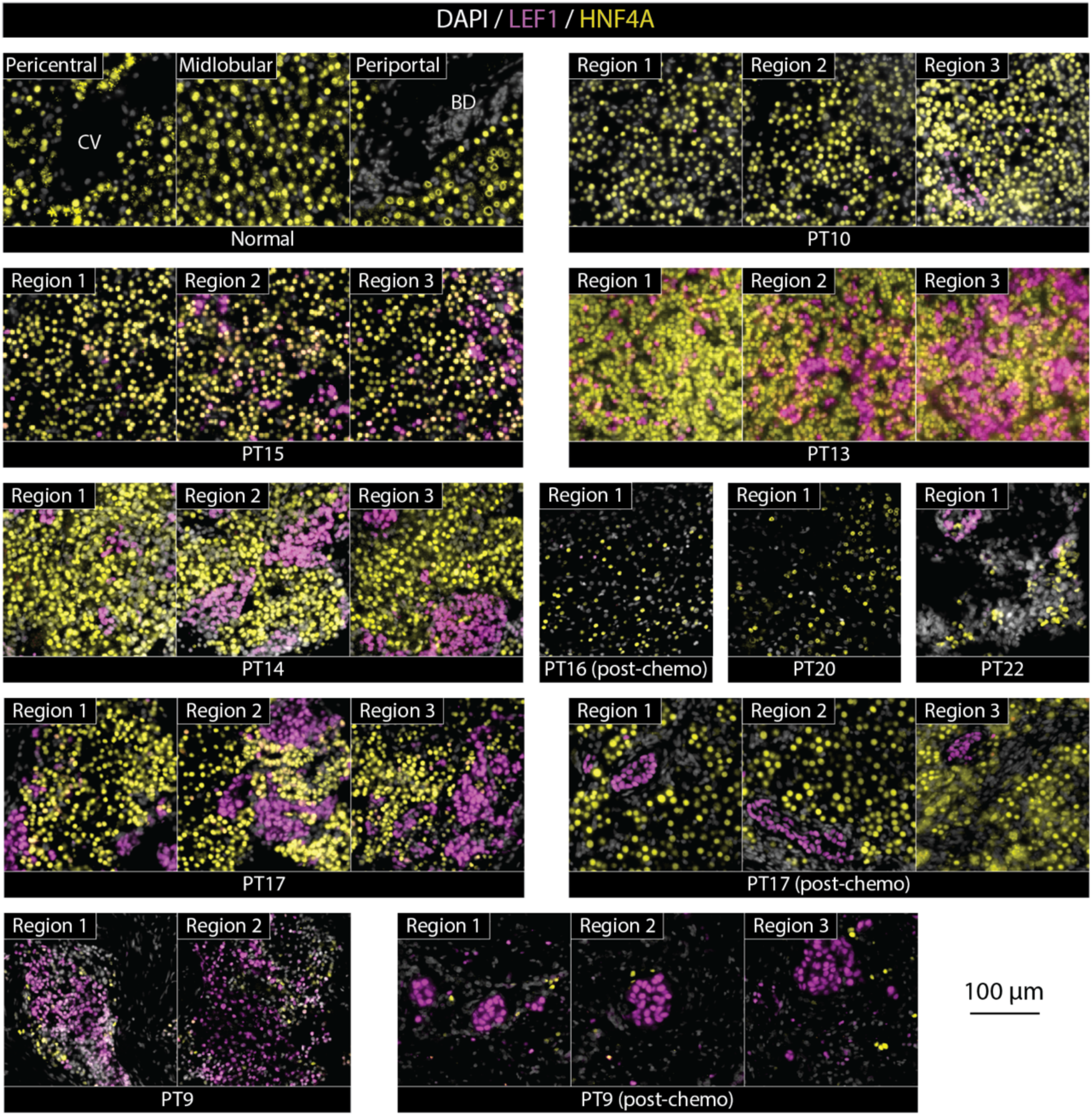
Immunofluorescence staining of TFs LEF1 and HNF4A mark distinct tumor subpopulations. Co-staining of HNF4A (yellow) and LEF1 (purple) in normal liver and tumor tissues displayed a mutually exclusive expression pattern, with multiple regions of the same tissue shown.

Previous studies have shown that embryonal hepatoblastoma regions correlate with diffuse nuclear and cytoplasmic β-catenin staining, while fetal regions correlate with moderate cytoplasmic and membranous staining along with focal nuclear staining^14,42^. Indeed, we found that in HNF4A^+^ cells, β-catenin is either localized to the membrane or shows diffuse cytoplasmic and membranous staining along with focal weak nuclear staining **(Figure S4)**. This is in contrast with LEF1^+^ cells where β-catenin shows strong nuclear staining but displays a higher degree of heterogeneity than LEF1 staining. In sum, we confirmed the existence of two distinct tumor subpopulations expressing either HNF4A or LEF1 and found that these two markers could distinguish tumor subpopulations better than β-catenin staining alone.

### Establishing patient-derived hepatoblastoma organoids from multiple clinical stages

The limited availability of pediatric liver tumor models that recapitulate the genomic and transcriptomic heterogeneity of the disease has hindered *in vitro* modeling. Herein, we established hepatoblastoma organoids using patient tumor material, obtained from biopsies or resected tumors. In total, we established a cohort of twelve tumor organoid models from ten patients, representing various clinical stages such as pre-chemotherapy, post-chemotherapy, relapse, and metastasis **(Figure 4A, Table S1)**. For two patients, PT13 and PT17, we established organoids from tumor material obtained at two different time-points: at diagnosis and during surgical resection.

**Figure 4.**
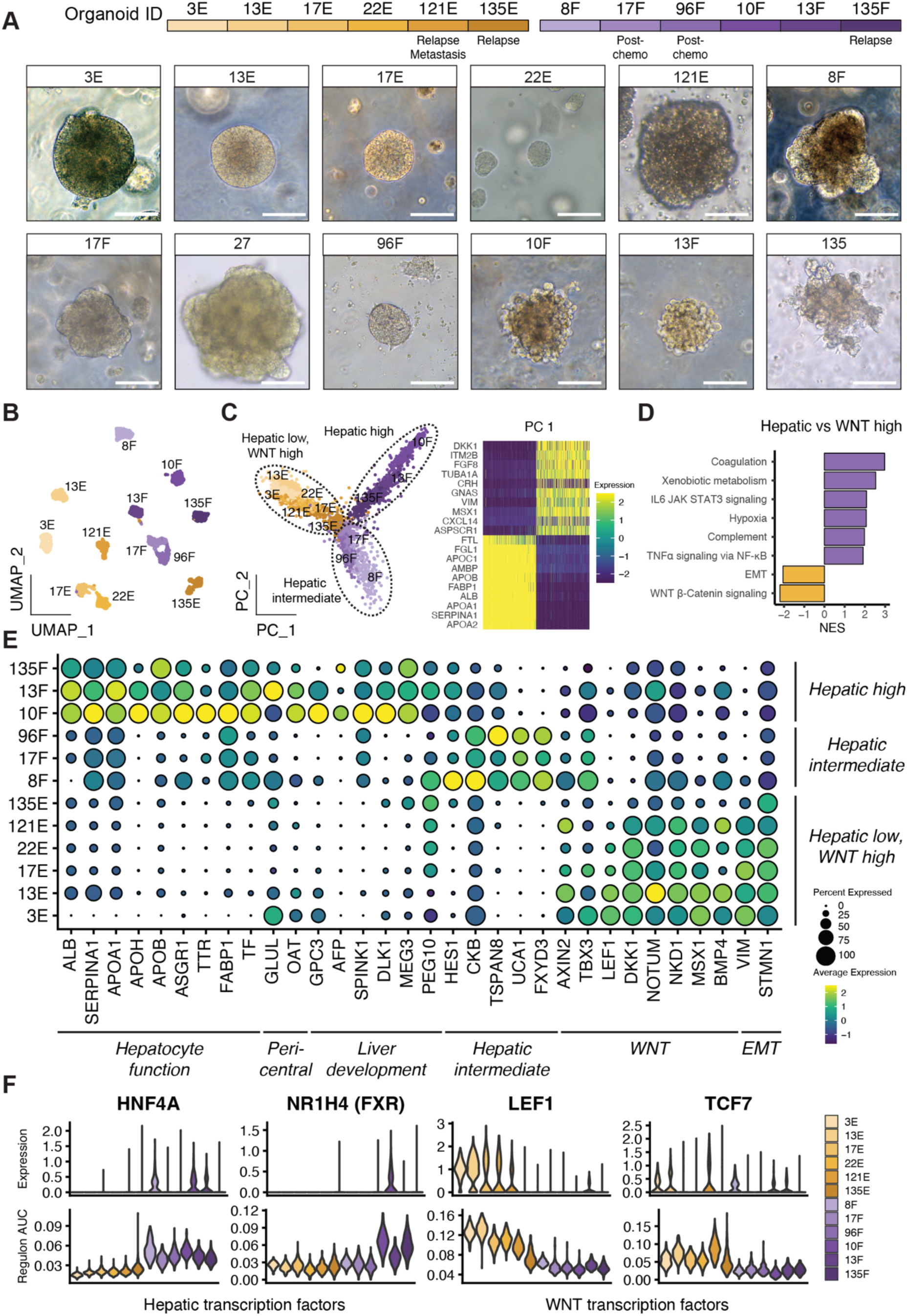
Single-cell RNA-seq analysis elucidates molecular heterogeneity of HB organoids. (A) Overview of different organoid lines and corresponding clinical stages (top) and phase-contrast images of organoids (bottom). Morphologically, embryonal-like organoids are more densely packed and have smooth surfaces while fetal-like organoids have more irregular shapes. Scale bars, 100 µm. (B) UMAP annotated by organoid model. (C) PCA plot showing three groups of organoids that can be distinguished based on the expression level of hepatic markers (left). Heatmap showing genes included in the first principal component (right). (D) Gene set enrichment analysis of hallmark gene sets comparing the ‘hepatic-high/intermediate’ groups with the ‘hepatic-low, WNT high’ organoids. (E) Dot plot showing expression of markers for the different groups of organoids. (F) Violin plots showing expression levels and regulon activity scores for hepatic transcription factors (HNF4A, NR1H4) and WNT/β-catenin transcription factors (LEF1, TCF7).

To confirm the tumor origin of the organoids, we performed targeted sequencing on *CTNNB1* exon 3 and verified the presence of point mutations or deletions, as identified in the original tumors, in all organoids **(Table S1**). Western blot analysis confirmed the reduction in β-catenin molecular weight in the organoids containing *CTNNB1* deletions. In addition, inferred copy number variation (CNV) profiles illustrated the presence of chromosomal aberrations **(Figure S5C, S5D).**

### Organoids recapitulate fetal- and embryonal-like tumor features

To obtain a transcriptomic landscape of the organoid cohort, we performed scRNA-seq **(Figure 4)**. Unsupervised graph-based clustering showed that organoids primarily clustered according to their sample origins (**Figure 4B**). However, principal component analysis (PCA) segregated organoids from various patients broadly into three groups, which we denoted as ‘hepatic-high’, ‘hepatic-intermediate’ and ‘hepatic-low, WNT-high’ based on marker expression (**Figure 4C-E)**. These included hepatic markers, such as *ALB*, *APOA1* and *SERPINA1* on the ‘hepatic-high’ group, and mesenchymal and WNT target genes, such as *DKK1*, *VIM* and *MSX1* on the ‘hepatic-low, WNT-high’ group.

Next, we compared the expression patterns across the organoid cohort **(Figure 4D, 4E, S5A)**. Hepatic TFs, hepatic functional markers (*e.g.,* coagulation, fatty acid metabolism), and fetal liver markers were expressed at varying levels in the ‘hepatic-high’ and ‘hepatic-intermediate’ groups but were absent or low in the ‘hepatic-low, WNT-high’ organoid group **(Figure 4D, 4E)**. These markers correspond with the fetal-like signature from the tumor tissue analyses. The ‘hepatic-low, WNT-high group’ was enriched with WNT target genes and EMT markers, including *AXIN2*, *LEF1*, *DKK1*, *NOTUM*, *NKD1* and *VIM*, in line with the embryonal-like tumor signature. In addition, the ‘hepatic-intermediate’ group (8F, 17F and 96F) expressed markers related to tumor progression and invasion, including *TSPAN8*, *CKB*, *UCA1* and *FXYD3* **(Figure 4E)**. However, two of the three samples (17F, 96F) were derived from post-chemotherapy tumor material and could only be cultured in a different medium (‘reduced medium’, see Methods and **Table S1**). Further studies are needed to investigate the biological implication of these observations.

We also performed gene regulatory network analysis on this dataset and identified WNT pathway regulons, including LEF1 and TCF7, specifically in the ‘hepatic-low, WNT-high’ organoids, while HNF4A was enriched in the hepatic tumor organoids **(Figure 4F, S5B)**. Another hepatic regulon, FXR, was enriched in the ‘hepatic-high’ cells but absent from the ‘hepatic-intermediate’ group **(Figure 4F)**. In summary, our analysis of the organoid cohort revealed that the expression patterns of hepatic markers, WNT pathway-related factors, and EMT markers aligned with the fetal- and embryonal-like hepatoblastoma signatures identified in the tumor tissue analyses. This suggests that our organoid models effectively recapitulate the characteristics of the distinct subpopulations observed in hepatoblastoma.

Next, to validate the gene regulatory network analysis, we performed single-cell Multiome ATAC and gene expression analysis on a subset of organoids **(Figure 5)**. The differential gene expression per sample is shown in **Figure S5E**. scATAC analysis identified an enrichment of TF motifs for hepatic development in the ‘hepatic-high/intermediate’ organoids, such as HNF family members (*HNF4A*, *4G*, *1B*, *1A*), *RXRG* and *PPARD* **(Figure 5A, 5B)**. Furthermore, the ‘hepatic-intermediate’ organoids showed enrichment of motifs of AP-1 family TFs (*FOS/JUN*), which are typically activated during liver regeneration^43^. The ‘hepatic low, WNT high’ tumor organoids were enriched with TF motifs related to the WNT pathway (*LEF1*, *TCF7*, *TCF7L2*), EMT (*MEOX2*) and apoptosis (*TP53*, *TP63, TP73*).

**Figure 5.**
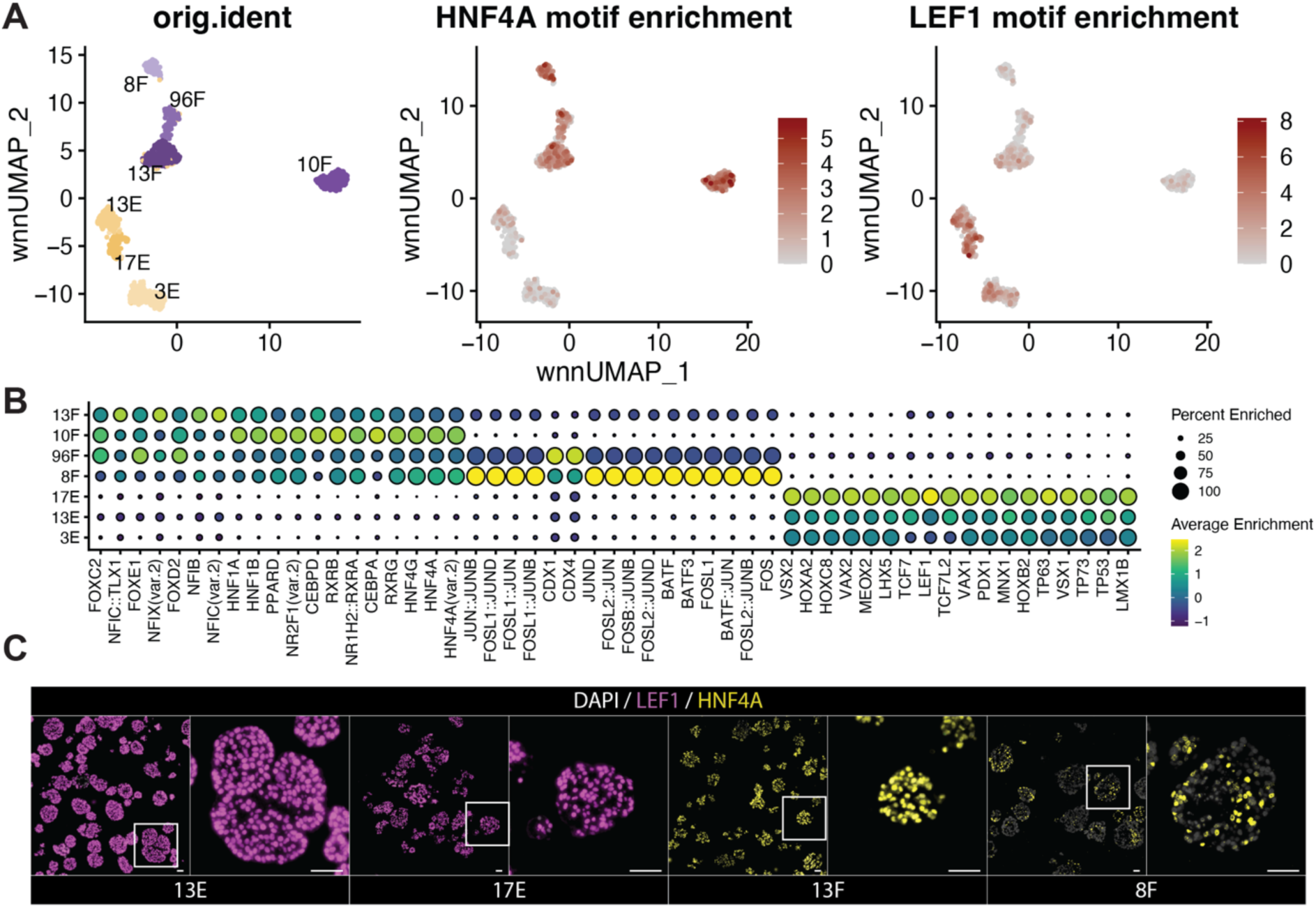
Single cell ATAC-seq identifies transcription factor motif enrichment associated with hepatoblastoma subtypes. (A) Weighted nearest neighbor (WNN) UMAP based on RNA and ATAC-seq data, separated by organoid model (left) and showing motif enrichment (right) for transcription factors HNF4A (fetal-like) and LEF1 (embryonal-like). (B) Dot plot showing the top enriched motifs per organoid model. (C) Immunofluorescence staining of FFPE sectioned organoid slides for LEF1 and HNF4A for embryonal lines 13E and 17E, and fetal lines 13F (hepatic-high) and 8F (hepatic-intermediate). Scale bars, 50 µm.

In sum, we were able to expand tumor organoids from the two major hepatoblastoma subtypes. The transcriptomic and chromatin accessibility landscapes of the organoids correspond with the fetal- and embryonal-like tumor signatures **(Figures 1-3)**. In addition, consistent with tumor tissue stainings, HNF4A and LEF1 marked fetal- and embryonal-like tumor organoids, respectively **(Figure 5C)**.

### Drug screening revealed distinct drug sensitivity profiles and potential therapeutic targets

To identify potential targeted treatment options against hepatoblastoma, we performed drug screening of over 200 compounds on 11 of our hepatoblastoma organoid models **(Figure 6, Table S1)**. To assess drug sensitivity, the area under the curve (AUC) for the dose-response curve of each compound was determined. These findings were depicted in a hierarchically clustered heatmap, providing insights into the drug sensitivity profiles, and highlighting correlations between the different organoid models (**Figure 6A, S6A**).

**Figure 6.**
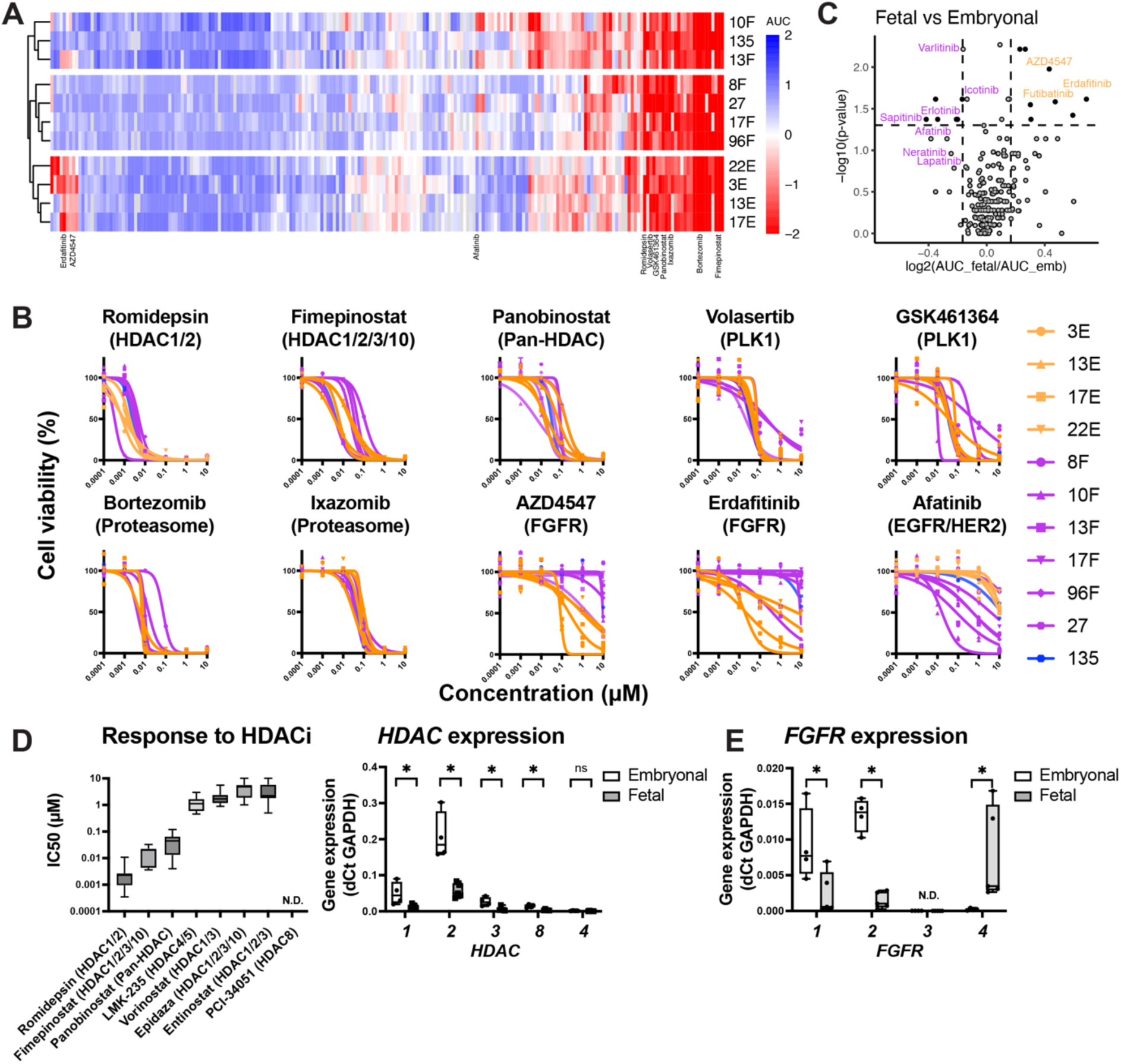
High-throughput drug screening of tumor organoids provides insight into targetable pathways. (A) Clustered heatmap, scaled by rows, of AUC values of the dose-response curves from the high-throughput drug screens of 11 organoid models. (B) Dose-response curves of selected drugs that are effective in all models, including those targeting HDAC, PLK1 and proteasomes, as well as drugs that show selective sensitivities in embryonal-like lines (targeting FGFR) and fetal-like lines (targeting EGFR). (C) Volcano plot revealing the most effective compounds specific for either the embryonal-(orange) and fetal-like (purple) tumor organoid models. (D) Boxplot showing IC50 values for all models for all HDAC inhibitors in the drug library (left). Expression of selected HDACs in organoid, measured by qRT-PCRs and separated by embryonal- and fetal-like models. Bars represent minimal and maximal values. (E) Boxplot showing expression of *FGFR* genes in organoids, measured by qRT-PCR and separated by embryonal- and fetal-like models. Bars represent minimal and maximal values.

We identified various classes of inhibitors targeting a wide range of hepatoblastoma organoids, including those that target HDAC, the proteasome, PLK-1 and FGFRs (**Figure 6B)**. Some of these compounds, such as vorinostat^44^ (an HDAC inhibitor), bortezomib^20^ (a proteasome inhibitor), and volasertib^45^ (a PLK-1 kinase inhibitor), were previously found to be effective against liver tumor cell lines and PDX models **(Figure 6A, 6B)**. However, compounds identified from other studies, such as olaparib^46^, a PARP1 inhibitor, were only moderately effective in our organoid models indicating potential biological differences between tumor cell lines and patient-derived organoid models (**Figure S6F**).

We further investigated the effect of different HDAC inhibitors on the organoid models. The organoids were sensitive to romidepsin (targeting HDAC1/2), panobinostat (pan HDAC), and fimepinostat (HDAC1/2/3/10), but not to entinostat (HDAC1/3) or PCI-34051 (HDAC8) treatment (**Figure 6B, S6D**). This finding indicates that only a subset of HDAC inhibitors may have therapeutic utility in hepatoblastoma, in agreement with a previous study^44^. The HDAC protein family consists of eleven HDACs, divided into four sub-classes based on sequence homologies. We assessed the mRNA expression of HDACs through scRNA-seq analysis, and observed the expression of *HDAC1*, *2* and to a lesser extent *HDAC 3* (**Figure S6B**).

We further assessed the mRNA expression of all Class I HDACs (*HDAC 1, 2, 3* and *8*), as well as previously reported *HDAC4 (*a Class IIa HDAC) in organoids using qRT-PCR (**Figure 6D, S6B**). *HDAC1* and *HDAC2* showed high expression levels across the cohort, with *HDAC2* expression being approximately 10-fold higher than that of *HDAC1*. The expression of *HDAC1/2* was also higher in embryonal-like subsets than fetal-like subsets (**Figure 6D**). *HDAC2* expression was also described previously by Cairo *et al.*^14^. Other HDACs, including *HDAC3*, *8* and *4* were expressed at lower levels than *HDAC1* and *2*. These data suggest that HDAC1 and 2 could be the possible therapeutic target of HDAC inhibitors in hepatoblastoma.

We then compared drug response profiles between fetal- and embryonal-like organoids **(Figure 7C)**. Particularly, fetal-like organoids were sensitive to treatment of small molecule inhibitors targeting the epidermal growth factor receptor (EGFR), such as afatinib, erlotinib and sapitinib, while embryonal-like organoids were sensitive to pan fibroblast growth factor receptor (FGFR) inhibitors, such as erdafitinib, futibatinib, and ponatinib (**Figure 6B, 6C, S6E**). scRNA-seq analysis revealed expression of FGFR1 and various FGFs, especially in the embryonal-like tumors and organoids (**Figure 6E, S6C**). We confirmed the mRNA expression of *FGFR1* and *FGFR2* in the embryonal-like organoids by qRT-PCR (**Figure 6E, S6C**), while *FGFR1* was highly expressed in embryonal-like tumor tissues (**Figure 1D**). This observation could indicate a previously unknown dependency on EGFR and FGFR signaling in the tumorigenesis of hepatoblastoma. We also noted the expression of *FGFR4* in the fetal-like organoids (Figure 6E), but not in fetal-like tumor tissues (**Figure 1D**). These organoids were not sensitive to pan-FGFR inhibitor treatment. Of note, FGF19, an FGFR4-specific ligand, was not included in our culture medium.

Finally, to identify compounds that specifically target hepatoblastoma, we compared the drug sensitivity profiles of hepatoblastoma organoids to a reference cohort consisting of other pediatric tumor model screens, which included Wilms tumor, malignant rhabdoid tumor, rhabdomyosarcoma, Ewing sarcoma, and neuroblastoma models^47,48^ (**Figure S7)**. We found that romidepsin was highly specific for hepatoblastoma compared to the reference cohort. Furthermore, we found that FGFR and EGFR inhibitors were highly specific for the respective hepatoblastoma subtypes compared to the reference cohort. In sum, our study identified drugs that could target both fetal and embryonal-like tumor organoids, as well as drugs that could selectively target a specific hepatoblastoma subtype.

## DISCUSSION

The majority of hepatoblastoma cases (> 90%) carry a mutation in the *CTNNB1* gene encoding for β-catenin, yet hepatoblastoma represents a disease with considerable molecular heterogeneity^2,49^. In this study, we analyzed patient tumor tissues and tumor organoids using scRNA-seq, ST, and scATAC-seq. Focusing on epithelial hepatoblastoma, we described two distinct tumor signatures. The first subpopulation, which we designated as ‘fetal-like’, expressed hepatic markers related to liver function, including pericentral markers typically activated by the WNT pathway, as well as previously described fetal liver markers^14^. Of note, in the normal liver, WNT signals, originating from endothelial cells of the central vein, play a crucial role in inducing the expression of pericentral genes in hepatocytes^50–52^.

In contrast, the second tumor subpopulation, which we annotated as WNT-high ‘embryonal-like’, displayed low levels of hepatic markers, but exhibited high levels of general WNT pathway target genes, including *AXIN2, APCDD1, NOTUM*, and *DKK1/4*. The fetal-like tumor signature is largely in agreement with previous bulk RNA-seq studies, which describe tumors enriched with hepatic features (termed ‘C1’^14^, ‘F’^16^, ‘C’^19^, ‘hepatocytic’^18^ or ‘hepatocyte’^17^), whereas the embryonal-like tumor signature is consistent with previously described tumors enriched with progenitor features (termed ‘C2A’^20,21^, ‘E’^16^, ‘A’^19^, ‘liver progenitor’^18^ or ‘proliferative’^17^). Our single-cell analysis further delineated the distinct tumor features discerned between the fetal- and embryonal-like tumor subgroups^13–19^.

A significant finding from our study is that, through gene regulatory network analysis, hepatic TFs such as HNF4A, FXR, CAR, PXR, and AR, are predominantly present in the fetal-like group but absent in the embryonal-like group. Conversely, embryonal-like tumors were enriched in key WNT pathway TFs, such as LEF1, TCF7, and TCF7L2. This finding was corroborated by the chromatin accessibility landscape analysis of a large cohort of tumor organoids established in this study. Furthermore, validation on tumor tissue sections demonstrated a mutually exclusive staining pattern of HNF4A and LEF1, in agreement with single-cell and spatial profiling. Taken together, our data suggests the presence of dichotomous expression patterns in hepatoblastoma, with intra- and inter-tumoral heterogeneity present within each tumor subset.

The absence of HNF4A in the embryonal-like tumor subset is intriguing, given that this TF is generally considered the central regulator of hepatic differentiation and is essential for maintaining liver function^41,53^. In previous studies, knockdown of *HNF4A* during hepatic development prohibited expression of many hepatic genes^54^. The absence of HNF4A, as well as other hepatic TFs, could explain the low levels of hepatic markers in the WNT-high embryonal tumor subset. Moreover, HNF4A has been previously shown to be essential for the transition of endoderm to a hepatic fate during development^55,56^. These observations suggest that hepatoblastoma could potentially arise at early stages of fetal liver developmental, with cellular differentiation being halted along the developmental trajectory.

While normal hepatocytes displayed predominantly membranous β-catenin staining, mutations in the *CTNNB1* gene in tumor cells resulted in the intracellular accumulation of β-catenin product. Previous studies^14,42^ have shown that embryonal regions correlate with diffuse nuclear and cytoplasmic β-catenin staining, while fetal regions correlate with moderate cytoplasmic and membranous staining along with focal nuclear staining. This is largely in agreement with our own observations. Based on the staining for HNF4A and LEF1, we were able to unequivocally distinguish these two hepatoblastoma subgroups, while it was not as evident with β-catenin staining alone. Currently, the classification of hepatoblastoma based on histology is complex due to the inherent molecular heterogeneity of the disease. This is further complicated by the presence of spatial heterogeneity, where fetal and embryonal tumor cells could be intermixed. Considering our finding, the use of LEF1 and HNF4A as biomarkers, in combination with existing markers, could enhance tumor molecular diagnostics, allowing for a more precise correlation between molecular profile and disease outcomes.

The distinct WNT signaling programs observed in the fetal- and embryonal-like tumor cells are noteworthy. In embryonal-like tumor cells, canonical WNT pathway target genes are abundant, consistent with prominent nuclear localization of β-catenin and presence of TCF/LEF family members. Separately, fetal-like cells displayed an enrichment of pericentral hepatic markers, typically activated by the WNT pathway, and a decrease in periportal hepatic markers repressed by the WNT pathway. A previous study focusing on liver zonation described the interaction between HNF4A, β-catenin and TCF7L2. Briefly, HNF4A can repress β-catenin-dependent transcription, and β-catenin can repress HNF4A-dependent transcription^57^. In line with this, a study using hepatocellular carcinoma (HCC) cells found that HNF4A competes with β-catenin for binding to TCF7L2 to facilitate repression of β-catenin/TCF7L2 target genes^58^. In addition, overexpression of HNF4A resulted in the depletion of nuclear β-catenin. The relationship between β-catenin and HNF4A in hepatoblastoma remains to be elucidated, although a similar process could be at play here. Separately, a recent study by Pagella *et al.*^38^ demonstrated that β-catenin binding to genomic loci is cell-type specific and associated with both activation and repression of gene expression programs, leading to divergent responses depending on the cellular context, and that β-catenin DNA binding is a highly dynamic and temporally regulated process.

Here, we established a large cohort of hepatoblastoma organoids from different clinical stages (pre-, post-chemotherapy, relapse) and performed drug screening using an extensive panel of compounds. We identified multiple compounds from various classes of inhibitors that could target a broad range of organoids, such as HDAC inhibitors, proteasome inhibitors, PLK-1 inhibitors and FGFR inhibitors. Of particular interest are HDAC inhibitors. We confirmed *HDAC1*/*2* expression in tumor organoids, with the embryonal tumors expressing higher levels of HDAC than the fetal tumors, which correlates with higher β-catenin activity observed in the embryonal subgroup. Histone deacetylase has been shown to function as a cofactor for β-catenin^59^. In the same study by Pagella *et al.*^38^, β-catenin was found to be dependent on HDAC for chromatin opening, and HDAC inhibition fully abolished its activity. In addition, many of these drugs identified in this study, including romidepsin and panobinostat, have been approved for clinical use^60–63^, presenting a great opportunity to assess their efficacy in pediatric liver cancer patients.

In addition, our study indicates, for the first time, the presence of distinct drug sensitivity profiles correlating with tumor heterogeneity. These profiles were characterized based on the tumor subtypes rather than clinical stage. The fetal-like tumor organoids were selectively targeted by EGFR inhibitors, and the embryonal-like organoids by FGFR inhibitors. This finding underscores the potential role of EGF and FGF pathways, in addition to the known role of the WNT pathway, in hepatoblastoma tumorigenesis. Relevant to this, a previous study identified *FGF8* as one of the most upregulated genes in metastatic hepatoblastoma^64^. These observed correlations could allow for the development of subtype-specific drug treatment or combination treatments with enhanced efficacy. Taken together, our findings may pave the way for improved hepatoblastoma treatment, offering hope for reduced mortality rates and enhanced quality of life in the future.

## ACKNOWLEDGEMENTS

We would like to extend our gratitude to the M4C Clinical team including Martine van Grotel, Kathelijne Kraal, Kees van de Ven, Lideke van der Steeg, Anneloes Bohte, Annemieke Littooij, Maarten Smits, Marijn Scheijde-Vermeulen, and to Liset Lansaat and Charlotte Aart. This work would not have been possible without support from the Princess Máxima Center Single-Cell Genomics facility (Tito Candelli and Aleksandra Balwierz), from the Big Data Core and Kemmeren group (Ianthe van Belzen and Jayne Hehir-Kwa), the Stunnenberg group (Zhijun Yu and Cristian Ruiz Moreno) and the Van Heesch group (Jip van Dinter).

## AUTHOR CONTRIBUTIONS

Conceptualization: W.C.P, J.Z. Methodology: W.C.P., T.A.K., Y.L., S.A.S., L.J.K. Investigation: T.A.K., Y.L., S.A.S., L.J.K., F.R., M.v.d.W., V.A., S.E. Software, analysis pipeline: P.L., W.L.M., T.M., J.D., T.A.K. Visualization: T.A.K, S.A.S., Y.L. Resources: W.C.P., H.C., J.Z., H.S., T.M., R.R.d.K., R.H.d.K., V.E.d.K, E.D., M.v.d.H., F.W.d.F., D.J.M., K.K. Supervision: W.C.P., H.C., H.S. Funding acquisition: W.C.P., H.C., J.Z., H.S., J.J.M., Writing: W.C.P., T.A.K., S.A.S. with inputs from all authors.

## METHODS

### Primary tumor tissues

Tumor material and distal normal liver tissue were obtained with written informed consent from biopsies and resections performed at the Princess Máxima Center for Pediatric Oncology Utrecht, the University Medical Center Groningen, and the Universitätsklinikum Münster. Patient samples and clinical data of the Dutch patients were obtained after approval by the Biobank and Data Access Committee of the PMC (PMCLAB2020-107).

### Organoid culture

Tissue dissociation was performed as described previously^22^. Briefly, tissues were minced into 1-2 mm pieces using small scissors in a glass Petri dish, followed by incubation in pre-warmed Liver Perfusion Medium (ThermoFisher) for 15 min at 37 °C. Next, minced tissues were washed once with DPBS and spun down at 300 x *g* for 3 minutes. The pellet was collected and incubated in pre-warmed digestion mix at 37 °C on a shaker at 150 rpm. Digestion mix consisted of Liver Digestion Medium (ThermoFisher) with 1% HEPES and 700U/mL Collagenase type IV (Worthington Biochemical). Dissociation progress was assessed after 15 and 30 minutes, and the samples were mechanically dissociated by pipetting up and down 10 times to obtain substantial dissociation. If necessary, red blood cell lysis buffer (Biolegend) was used, following manufacturer’s instructions. Finally, the enzyme was washed off using the Hepatocyte Wash Medium (ThermoFisher) and the remaining cells or small aggregates were mixed with 100 % reduced growth factor basement membrane extract (BME; R&D) and plated in 20 µL droplets on culture plate that had been incubated overnight. Occasionally, no digestion was necessary, and aggregates were plated immediately after mincing. Organoids were cultured in culture medium described previously, with slight modifications^65–67^. The final culture medium (termed ‘full medium’) consisted of: Advanced DMEM/F12 (Invitrogen) containing HEPES, GlutaMAX, Penicillin-Streptomycin and Normocin, supplemented with 2 % B27 (Gibco), 1 % N2 (Gibco), 1.25 mM N-Acetylcysteine (Sigma), 10 nM gastrin (Sigma), 50 ng/mL EGF (Peprotech), 10% RSPO1 conditioned media (produced in house), 100 ng/mL FGF10 (Peprotech), 25 ng/mL HGF (Peprotech), 10 mM Nicotinamide (Sigma), 5 µM A83.01 (Tocris), 10 µM forskolin (Tocris) and 10 µM Y27632 (Sigma Aldrich) and 0.5 nM next-generation WNT surrogate^68^. In later established models, FGF10 was omitted and did not affect organoid establishment or growth (for the drug screens, this was only included in 10F, 13E, 13F, 17E and 17F). Several lines (17F, 96F, 27) could not be expanded in the ‘full medium’ without forming cystic organoids typically observed in cholangiocyte organoid culture. For these lines, we used the medium as described in Wu *et al.*^66^. Compared to the ‘full medium’, the reduced medium lacks gastrin, forskolin, RSPO1 and WNT, and is supplemented with 3 nM dexamethasone (Bio-Techne). Organoids could be passaged every 1-3 weeks and varied between tumors. Passaging was performed as described before^69^. Briefly, BME was first dissociated using dispase (STEMCELL Technologies), before washing at least twice with PBS supplemented with 5 % FBS (v/v). Organoids were then incubated in TryPLE Express (Gibco) at 37 °C and pipetted to obtain small aggregates of cells and washed once more before replating in BME.

### Single-cell RNA and ATAC sequencing

Single-cell analysis of tissue and organoid samples was performed using the 10x Genomics Single-Cell Expression platforms according to the manufacturer’s protocols (Chromium Next GEM Single-Cell 3’ Reagent Kits v3.1 and Chromium Next GEM Single-Cell Multiome ATAC + Gene Expression). In brief, fresh tissues were minced (<2 mm^2^) and viably frozen until processed. For scRNA-seq, minced tissue was rapidly defrosted, washed and dissociated for 1 hour at 37 °C, 250 rpm using dissociation mix 1. Dissociation mix 1 contained 0.5 mg/mL Liberase (ThermoFisher) and 1 mg/mL Collagenase type IV (Gibco) in DMEM/F12 supplemented with Glutamax and 20 U/mL DNase (ThermoFisher). Red blood cells and dead cells were removed using an RBC lysis buffer (BioLegend) and a Dead Cell Removal kit (Miltenyi Biotec), respectively. Prior to loading, the cells were passed through Flowmi cell strainers (40 μm, Merck) and counted using a Bürker chamber.

For the scRNA-seq of the organoids, 3E and 8F organoids were sequenced individually, while the other lines were pooled together using the 3’ CellPlex Multiplexing Kit. In brief, organoids were dissociated using TryPLE Express into single cells and incubated with a unique 10x CellPlex molecular tag for 15 min at RT while shacking (250 rpm) before washing and pooling. Demultiplexing was performed during data analysis using Cell Ranger and all lines were integrated and analyzed together. For the organoid Multiome samples, organoids were first dissociated into single cells using TryPLE Express. Nuclei were isolated with NP40 lysis buffer with a 5 min incubation on ice and passed through a 70 μm cell strainer (Greiner Bio-One EASYstrainer, ThermoFisher). Samples were pooled based on nuclei concentration (Countess II cell counter, ThermoFisher) and sorted on a Sony SH800S cell sorter with a 100 μm nozzle for 7AAD (Invitrogen) positive singlets. After nuclei permeabilization samples were counted (Countess II). SNP-based demultiplexing was performed using Python packages cellsnp lite and Vireo. Genotyping references of the donors was obtained from whole-genome sequencing or whole-exome sequencing data of the tumor biopsy samples generated in the diagnostic setting. We recovered on average approximately 2200 and 1200 cells per organoid line for individually sequenced and multiplexed samples, respectively.

### Spatial RNA sequencing

Fresh tissues were snap frozen in isopentane (Sigma) chilled by liquid nitrogen. Frozen tissue pieces were embedded in Tissue-Tek O.C.T. Compound (Sakura) and stored at -80 °C until cryosectioning. Tissues were selected based on tissue histology and RNA quality (RIN scores > 2.4 of TRIzol isolated RNA). ST was performed using the Visium Spatial Gene Expression Solution (10x Genomics) according to the manufacturer’s protocols. In brief, 10 μm thick tissue sections were cut in a cryostat (Cryostar NX70 Thermo Fisher) and placed within the capture area of Visium Spatial Gene Expression Slides. Tissues were fixed in chilled methanol and H&E stainings were performed to assess tissue morphology and quality. The slides were imaged using a brightfield microscope (Leica DMi8 S platform). Tissue permeabilization times for normal liver and tumor tissue was optimized using the Visium Spatial Tissue Optimization workflow. Permeabilization times were set at 12 minutes for tumor tissue and 18 minutes for normal liver tissue. Full length cDNA and libraries were analyzed using a Qubit 4 fluorometer and an Agilent 2100 Bioanalyzer. cDNA libraries were sequenced on a NovaSeq6000 System (Illumina) with sequencing settings recommended by 10x Genomics.

### Single-cell RNA data analysis

Single-cell transcriptome data of hepatoblastoma and paired normal liver were obtained from Song *et al.*^22^ and accessed through the Gene Expression Omnibus (GEO) database (accession number GSE186975). Raw UMI-collapsed read-count data was analyzed through Seurat^70^ (version 4.3.0) in R (version 4.1.2). First, contamination by ambient RNA was estimated and removed by DecontX^71^ (celda version 1.10.0) using default settings. Next, low-quality cells were filtered by removing cells with fewer than 500 genes or a percentage of mitochondrial genes greater than 20%. Additionally, a maximum threshold for number of genes was set for individual samples. Separate workflows were implemented for dimensionality reduction and gene expression analyses. For dimensionality reduction and clustering, normalization was performed by SCTransform (version 0.3.5), and cell-cycle and related genes were identified as previously described^72^. We then integrated the data from different batches using fastMNN batch correction (SeuratWrappers version 0.3.1), while removing the cell cycle and related genes from the integration features. Dimensionality reduction and clustering were performed on the fastMNN-corrected data, using 40 dimensions and a resolution of 0.5, yielding 22 clusters. For gene expression analyses, counts were normalized and log-transformed by Seurat’s “LogNormalize” method, followed by identification of the 2000 most variably expressed genes and linear transformation. Differentially expressed genes (DEG) for each cluster were identified by Seurat’s FindAllMarkers and three genes with the highest fold-change expression were selected for each cluster to facilitate cell type identification. All clusters containing tumor cells or epithelial cells of the normal liver were selected (clusters 4, 8, 11, 13 and 18) and reprocessed by the workflow described above to further identify epithelial and tumor subpopulations. A cluster of tumor cells expressing neuroendocrine markers, which was described in Song *et al.*, was specific to only one patient. Therefore, we did not focus on this cluster and excluded it for this analysis. In addition, because some of the identified clusters contained low quality cells, contaminated by ambient RNA or cell fragments, high quality clusters were selected (clusters 2, 3, 5, 6 and 8) and reprocessed once more by the same workflow, creating the final subset comprising 4 clusters.

For the primary tumor from PT13 and the organoid lines, raw reads were aligned using CellRanger, omitting intronic reads. The Seurat package standard workflow was employed as described before. Low-quality cells and doublets were filtered out from the primary cells and organoids separately based on number of unique genes measured and the percentage of mitochondrial genes. After filtering, data was processed using the NormalizeData, ScaleData and FindVariableFeatures functions for the RNA assay, and SCTransform to create an SCT assay. Dimensionality reduction was performed on the SCT assay using RunPCA, and UMAP coordinates were calculated using RunUMAP. Confounding genes were removed from the VariableFeatures as described above. Two separate clusters of organoid sample 135 were identified and labeled 135F and 135E. All organoid idents were subsetted to 269 cells each. Dot plots and heatmaps were made using DoHeatmap or DotPlot, or the pheatmap package, after markers were calculated using the FindAllMarkers or FindMarkers commands, using the RNA assay. The hierarchical clustering plot was made using BuildClusterTree. InferCNV was run using standard settings, with hepatocytes from the tissue object as reference^73^. pySCENIC was run on a high-performance cluster with standard settings, using a singularity file obtained from the Aerts lab^31^. After obtaining the AUC scores, Seurat was used to calculate differentially active regulons, using FindAllMarkers with logfc.threshold = 0.005. Gene set enrichment analysis was run using the fgsea package (v 1.24.0), using h.all.v7.4.symbols.gmt.

### Spatial data analysis

Raw FASTQ files and histology images were processed using Space Ranger software (v 1.2.2). Each sample was normalized individually using the SCTransform function of the Seurat R package with default parameters, except method = “glmGamPoi” to improve the speed, and return.only.var.genes = FALSE. Clustering was performed using FindNeighbors and FindClusters. Clusters were annotated based on marker gene expression, differential gene expression using FindAllMarkers and tissue histology. Clusters with similar gene expression profiles were combined. All samples were then merged with the merge function of the Seurat R package with default parameters.

### Organoid multiome analysis

Raw reads were aligned using CellRanger. Joint analysis was performed using the Seurat and Signac packages^74,75^, following standard workflow unless specified otherwise. Cells were filtered using the following settings: “nCount_ATAC < 40000 & nCount_ATAC > 100 & nCount_RNA < 8000 & nCount_RNA > 300 & percent.mt < 2 & TSS.enrichment > 3 & nucleosome_signal < 2”. For processing of the RNA assay, again SCTransform was used, mitochondrial and cell cycle (correlated) genes were removed from the VariableFeatures as before, and principal component analysis was performed using the RunPCA command. For the ATAC assay, peak calling was performed using macs2. Dimensionality reduction was then performed by running RunTFIDF, FindTopFeatures and RunSVD. Next, multimodal analysis of both assays was performed, running FindMultiModalNeighbors with reduction.list = list(“pca”, “lsi”) and dims.list = list(1:50, 2:50); RunUMAP with nn.name = “weighted.nn” and reduction.name = “wnn.umap“; and FindClusters with graph.name = “wsnn”, algorithm = 3, and resolution = 0.2. Different lines were separated based on SNP based demultiplexing and marker expression. For downstream analysis, all lines were downsampled to 400 cells each. For motif analysis, ChromVar was run after adding the human JASPAR2020 motifs. Differentially enriched motifs were calculated using wilcoxauc and plotted.

### Immunofluorescence staining

Whole pieces of tissues were fixed in 10% Neutral Buffered Formalin for 1 hour to overnight, depending on the size of the specimen. Fixed tissues were processed using the Excelsior™ AS Tissue Processor (Thermo Fisher) following a standard protocol and embedded in paraffin blocks. Organoids were released from the BME using dispase, washed, and fixed for 1 hour in 10% Neutral Buffered Formalin. The FFPE blocks were sectioned into 4 µm thick sections that were placed on SuperFrost Plus slides (Thermo Fisher). Tissues sections were deparaffinized and rehydrated by immersing the slides in 3 changes of xylene followed by a series of decreasing concentrations of ethanol and rinsing in distilled water. Heat-induced antigen retrieval was performed by placing the slides in an appropriate buffer (see below) and boiling for 20 min using a double boiler method. For IF staining, tissue sections were permeabilized in 0.3% Triton X-100 in PBS (PBS-Tx) for 5-10 min. Next, blocking buffer (5% normal donkey serum [Jackson ImmunoResearch] in 0.1% PBS-Tx) was applied for 1 hour at room temperature (RT). Tissue sections were incubated with primary antibodies (see below) diluted in blocking buffer for 1 hour at RT or at 4 °C overnight and washed with 3 changes of 0.1% PBS-Tx. Secondary antibodies were diluted in PBS and applied for 1 hour at RT. All secondary antibodies were raised in donkey and conjugated to Alexa Fluor dyes (488, 555 and 647) (Thermo Fisher). Sections were washed with changes of PBS and counterstained with DAPI (Sigma Aldrich) at 5 µg/ml for 5 min at RT. Slides were mounted in 80% glycerol in PBS and a #1.5 coverslip. Images were acquired with 20X dry and 40X water immersion objectives on a Leica DMi8 Thunder widefield microscope equipped with 4 LED light sources (DAPI [395/25], FITC [475/28], Cy3 [555/28] and Cy5 [635/22]). Acquired images were adjusted for brightness and pseudo colored using FIJI software.

| Target | Supplier | Antigen retrieval | Dilution |
| --- | --- | --- | --- |
| HNF4A | MA1-199 (Thermo Fisher) | Citrate/tris | 1:50 |
| β-catenin | 610154 (BD Bioscience) | Citrate/tris | 1:100 |
| LEF1 | 2230 (Cell Signaling Tech) | Citrate/tris | 1:100 |

### High-throughput drug screening

Organoids were recovered from their BME matrix by dispase incubation, followed by three washing steps with cold PBS containing 5% FBS. Organoids were then resuspended in organoid medium supplemented with 5% BME and plated in Corning 384 well microplates (Sigma, CLS3764) at ±50 organoids per well, using a Multidrop Combi Reagent Dispenser (Thermo Scientific). The next day, cell viabilities of one plate were measured by a Spectramax i3x plate reader using the CellTiter-Glo 3D Cell Viability Assay (CTG3D), and drugs were added to the other plates, using an Echo 550 liquid handler. We used the PMC library (in-house developed library with >200 drugs specific for pediatric cancer). Each drug was tested in duplicate and in 6 different concentrations (0.1 nM to 10 μM in increments of factor 10). After 120 hours, viabilities of all wells were measured using CTG3D. Graphs were then fitted and area under the curve scores (AUCs) calculated using R. A volcano plot was generated using the Wilcoxon signed-rank test and the average AUC values for the two sub-groups based on clustering. Z-scores against the pediatric tumor reference cohort were calculated using (x - μ) / σ, where x represents the AUC of each drug in hepatoblastoma organoids, and μ and σ are the mean and standard deviation of the AUC values for that drug in other pediatric tumors, respectively. Drugs with IC50 values higher than 10 mM were excluded from the analysis.

### Sanger sequencing

Genomic DNA was extracted using the DNeasy Blood & Tissue Kit (Qiagen) and *CTNNB1* exon 3 was amplified using GoTaqβ G2 Flexi DNA Polymerase (Promega) according to the manufacturer’s protocol. The PCR product and sequencing primers were sent to Macrogen Europe to perform Sanger sequencing. The sequences of the PCR and sequencing primers were:

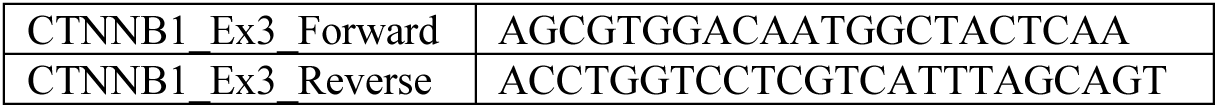

### RNA isolation and qRT-PCR

Total RNA was extracted using the RNeasy Mini Kit (Qiagen) and reverse transcribed using the GoScript™ Reverse Transcriptase (Promega) according to the manufacturer’s protocol. Quantitative RT-PCR was performed with GoTaqβ qPCR and RT-qPCR Systems (Promega) on a CFX384 Real-time System (Bio-Rad). Relative target gene expression levels were calculated using the delta-delta CT method. The primers were all ordered from Integrated DNA Technologies (IDT), with sequences shown below:

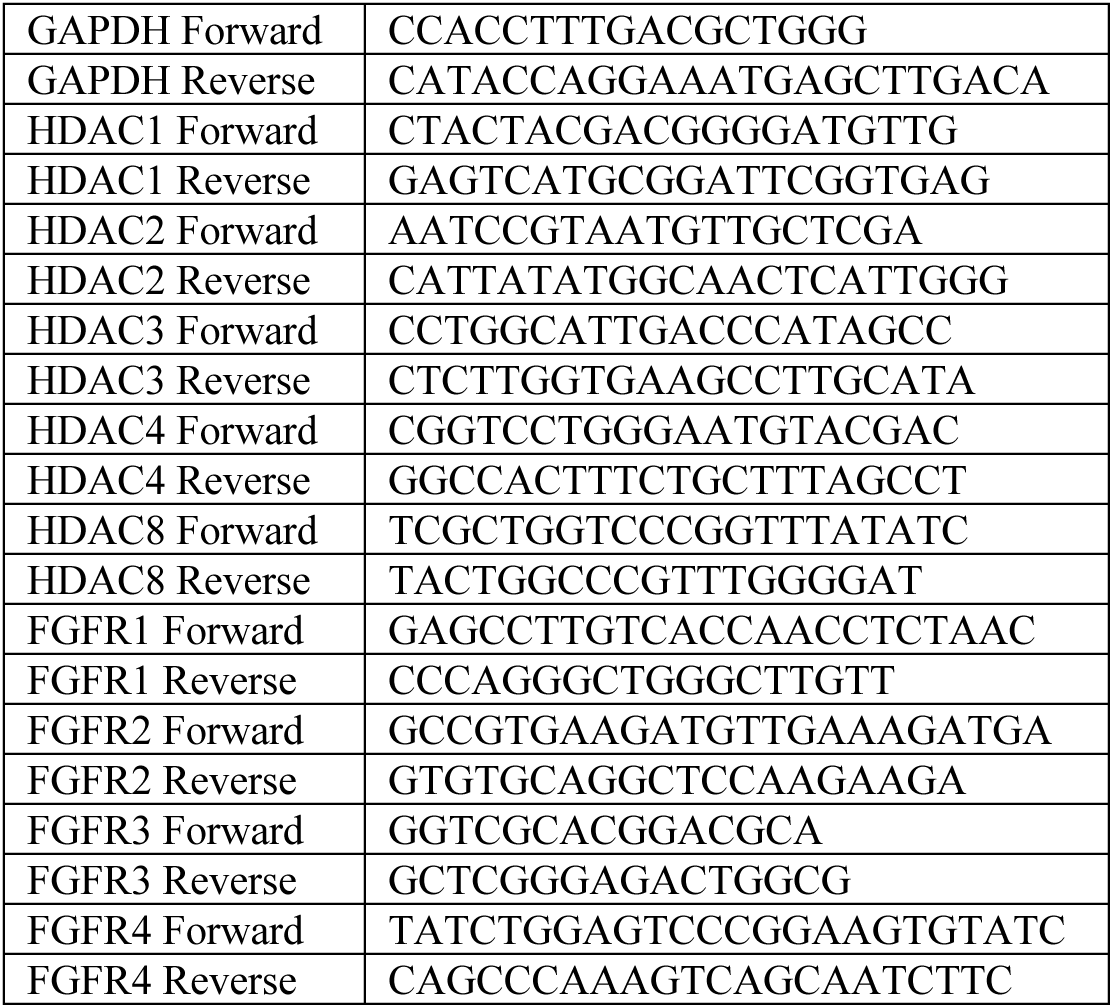

### Western blot

Cells were lysed in homemade lysis buffer. After quantification using a Qubit protein assay kit (Invitrogen), protein samples were separated by sodium dodecyl sulfate-polyacrylamide gel electrophoresis (SDS-PAGE). These were then transferred onto polyvinylidene difluoride (PVDF) membranes (Millipore). Membranes were blocked in 5% BSA (Thermo Fisher Scientific) for 1h at room temperature and incubated with antibodies against β-catenin (1:1,000; CST), followed by incubation with anti-rabbit (1:5,000) secondary antibodies conjugated to fluorescent dyes (IRDye). Immunoreactive proteins were subsequently visualized using the Odyssey CLx Infrared Imaging System.

### *CTNNB1* mutation analysis

Sequencing analysis of tumor biopsy materials was performed in the diagnostic setting using the institute’s standardized analysis pipelines. Single nucleotide variants and small indels in the *CTNNB1* gene (ENST00000349496.11/ ENSP00000344456.5) were assessed using whole-exome and/or whole-genome sequencing. Exon 3 deletions were assessed based on bulk RNA-sequencing analysis after realignment of the BAM files using STAR aligner (v2.7.8a).

## REFERENCES

1. Meyers, R.L., Aronson, D.C., and Zimmermann, A. (2012). Pediatric Surgery (Seventh Edition). 463–482. 10.1016/b978-0-323-07255-7.00033-7.

2. Ranganathan, S., Lopez-Terrada, D., and Alaggio, R. (2020). Hepatoblastoma and Pediatric Hepatocellular Carcinoma: An Update. Pediatr Devel Pathol 23, 79–95. 10.1177/1093526619875228.

3. Dasgupta, P., Henshaw, C., Youlden, D.R., Aitken, J.F., Sullivan, A., Irving, H., and Baade, P.D. (2020). Global trends in incidence rates of childhood liver cancers: A systematic review and meta-analysis. Paediatr. Périnat. Epidemiology 34, 609–617. 10.1111/ppe.12671.

4. Zsiros, J., Brugieres, L., Brock, P., Roebuck, D., Maibach, R., Zimmermann, A., Childs, M., Pariente, D., Laithier, V., Otte, J.-B., et al. (2013). Dose-dense cisplatin-based chemotherapy and surgery for children with high-risk hepatoblastoma (SIOPEL-4): a prospective, single-arm, feasibility study. Lancet Oncol 14, 834–842. 10.1016/s1470-2045(13)70272-9.

5. Perilongo, G., Maibach, R., Shafford, E., Brugieres, L., Brock, P., Morland, B., Camargo, B. de, Zsiros, J., Roebuck, D., Zimmermann, A., et al. (2009). Cisplatin versus Cisplatin plus Doxorubicin for Standard-Risk Hepatoblastoma. New Engl J Medicine 361, 1662–1670. 10.1056/nejmoa0810613.

6. Zsíros, J., Maibach, R., Shafford, E., Brugieres, L., Brock, P., Czauderna, P., Roebuck, D., Childs, M., Zimmermann, A., Laithier, V., et al. (2010). Successful Treatment of Childhood High-Risk Hepatoblastoma With Dose-Intensive Multiagent Chemotherapy and Surgery: Final Results of the SIOPEL-3HR Study. J Clin Oncol 28, 2584–2590. 10.1200/jco.2009.22.4857.

7. Moroz, V., Morland, B., Tiao, G., Hiyama, E., Kearns, P., and Wheatley, K. (2015). The paediatric hepatic international tumour trial (PHITT): clinical trial design in rare disease. Trials 16, P224. 10.1186/1745-6215-16-s2-p224.

8. Dembowska-Bagińska, B., Więckowska, J., Brożyna, A., Święszkowska, E., Ismail, H., Broniszczak-Czyszek, D., Stefanowicz, M., Grajkowska, W., and Kaliciński, P. (2019). Health Status in Long-Term Survivors of Hepatoblastoma. Cancers 11, 1777. 10.3390/cancers11111777.

9. Gröbner, S.N., Worst, B.C., Weischenfeldt, J., Buchhalter, I., Kleinheinz, K., Rudneva, V.A., Johann, P.D., Balasubramanian, G.P., Segura-Wang, M., Brabetz, S., et al. (2018). The landscape of genomic alterations across childhood cancers. Nature 555, 321–327. 10.1038/nature25480.

10. Eichenmüller, M., Trippel, F., Kreuder, M., Beck, A., Schwarzmayr, T., Häberle, B., Cairo, S., Leuschner, I., Schweinitz, D. von, Strom, T.M., et al. (2014). The genomic landscape of hepatoblastoma and their progenies with HCC-like features. J Hepatol 61, 1312– 1320. 10.1016/j.jhep.2014.08.009.

11. López-Terrada, D., Gunaratne, P.H., Adesina, A.M., Pulliam, J., Hoang, D.M., Nguyen, Y., Mistretta, T.-A., Margolin, J., and Finegold, M.J. (2009). Histologic subtypes of hepatoblastoma are characterized by differential canonical Wnt and Notch pathway activation in DLK+ precursors. Hum Pathol 40, 783–794. 10.1016/j.humpath.2008.07.022.

12. Vasudevan, S.A., Meyers, R.L., Finegold, M.J., López-Terrada, D., Ranganathan, S., Dunn, S.P., Langham, M.R., McGahren, E.D., Tiao, G.M., Weldon, C.B., et al. (2022). Outcomes of children with well-differentiated fetal hepatoblastoma treated with surgery only: Report from Children’s Oncology Group Trial, AHEP0731. J. Pediatr. Surg. 57, 251–256. 10.1016/j.jpedsurg.2022.05.022.

13. Hiyama, E. Gene expression profiling in hepatoblastoma cases of the Japanese Study Group for Pediatric Liver Tumors-2 (JPLT-2) trial. 10.31487/j.ejmc.2018.01.003.

14. Cairo, S., Armengol, C., Reyniès, A.D., Wei, Y., Thomas, E., Renard, C.-A., Goga, A., Balakrishnan, A., Semeraro, M., Gresh, L., et al. (2008). Hepatic Stem-like Phenotype and Interplay of Wnt/β-Catenin and Myc Signaling in Aggressive Childhood Liver Cancer. Cancer Cell 14, 471–484. 10.1016/j.ccr.2008.11.002.

15. Sumazin, P., Chen, Y., Treviño, L.R., Sarabia, S.F., Hampton, O.A., Patel, K., Mistretta, T., Zorman, B., Thompson, P., Heczey, A., et al. (2017). Genomic analysis of hepatoblastoma identifies distinct molecular and prognostic subgroups. Hepatology 65, 104–121. 10.1002/hep.28888.

16. Sekiguchi, M., Seki, M., Kawai, T., Yoshida, K., Yoshida, M., Isobe, T., Hoshino, N., Shirai, R., Tanaka, M., Souzaki, R., et al. (2020). Integrated multiomics analysis of hepatoblastoma unravels its heterogeneity and provides novel druggable targets. Npj Precis Oncol 4, 20. 10.1038/s41698-020-0125-y.

17. Nagae, G., Yamamoto, S., Fujita, M., Fujita, T., Nonaka, A., Umeda, T., Fukuda, S., Tatsuno, K., Maejima, K., Hayashi, A., et al. (2021). Genetic and epigenetic basis of hepatoblastoma diversity. Nat Commun 12, 5423. 10.1038/s41467-021-25430-9.

18. Hirsch, T.Z., Pilet, J., Morcrette, G., Roehrig, A., Monteiro, B.J., Molina, L., Bayard, Q., Trepo, E., Meunier, L., Caruso, S., et al. (2021). Integrated genomic analysis identifies driver genes and cisplatin-resistant progenitor phenotype in pediatric liver cancer. Cancer Discov 11, candisc.1809.2020. 10.1158/2159-8290.cd-20-1809.

19. Huang, H., Wu, L., Lu, L., Zhang, Z., Qiu, B., Mo, J., Luo, Y., Xi, Z., Feng, M., Wan, P., et al. (2022). Single-cell Transcriptomics Uncovers Cellular Architecture and Developmental Trajectories in Hepatoblastoma. Hepatology. 10.1002/hep.32775.

20. Hooks, K.B., Audoux, J., Fazli, H., Lesjean, S., Ernault, T., Dugot-Senant, N., Leste-Lasserre, T., Hagedorn, M., Rousseau, B., Danet, C., et al. (2018). New insights into diagnosis and therapeutic options for proliferative hepatoblastoma. Hepatology 68, 89–102. 10.1002/hep.29672.

21. Carrillo-Reixach, J., Torrens, L., Simon-Coma, M., Royo, L., Domingo-Sàbat, M., Abril-Fornaguera, J., Akers, N., Sala, M., Ragull, S., Arnal, M., et al. (2020). Epigenetic footprint enables molecular risk stratification of hepatoblastoma with clinical implications. J Hepatol 73, 328–341. 10.1016/j.jhep.2020.03.025.

22. Song, H., Bucher, S., Rosenberg, K., Tsui, M., Burhan, D., Hoffman, D., Cho, S.-J., Rangaswami, A., Breese, M., Leung, S., et al. (2022). Single-cell analysis of hepatoblastoma identifies tumor signatures that predict chemotherapy susceptibility using patient-specific tumor spheroids. Nat Commun 13, 4878. 10.1038/s41467-022-32473-z.

23. Bondoc, A., Glaser, K., Jin, K., Lake, C., Cairo, S., Geller, J., Tiao, G., and Aronow, B. (2021). Identification of distinct tumor cell populations and key genetic mechanisms through single cell sequencing in hepatoblastoma. Commun Biology 4, 1049. 10.1038/s42003-021-02562-8.

24. Benhamouche, S., Decaens, T., Godard, C., Chambrey, R., Rickman, D.S., Moinard, C., Vasseur-Cognet, M., Kuo, C.J., Kahn, A., Perret, C., et al. (2006). Apc Tumor Suppressor Gene Is the “Zonation-Keeper” of Mouse Liver. Dev Cell 10, 759–770. 10.1016/j.devcel.2006.03.015.

25. Lotto, J., Drissler, S., Cullum, R., Wei, W., Setty, M., Bell, E.M., Boutet, S.C., Nowotschin, S., Kuo, Y.-Y., Garg, V., et al. (2020). Single-Cell Transcriptomics Reveals Early Emergence of Liver Parenchymal and Non-parenchymal Cell Lineages. Cell 183, 702–716.e14. 10.1016/j.cell.2020.09.012.

26. Wesley, B.T., Ross, A.D.B., Muraro, D., Miao, Z., Saxton, S., Tomaz, R.A., Morell, C.M., Ridley, K., Zacharis, E.D., Petrus-Reurer, S., et al. (2022). Single-cell atlas of human liver development reveals pathways directing hepatic cell fates. Nat Cell Biol, 1–12. 10.1038/s41556-022-00989-7.

27. Cavard, C., Terris, B., Grimber, G., Christa, L., Audard, V., Radenen-Bussiere, B., Simon, M.-T., Renard, C.-A., Buendia, M.-A., and Perret, C. (2006). Overexpression of regenerating islet-derived 1 alpha and 3 alpha genes in human primary liver tumors with β-catenin mutations. Oncogene 25, 599–608. 10.1038/sj.onc.1208860.

28. Grozdanov, P.N., Yovchev, M.I., and Dabeva, M.D. (2006). The oncofetal protein glypican-3 is a novel marker of hepatic progenitor/oval cells. Lab. Investig. 86, 1272–1284. 10.1038/labinvest.3700479.

29. Shimomura, Y., Agalliu, D., Vonica, A., Luria, V., Wajid, M., Baumer, A., Belli, S., Petukhova, L., Schinzel, A., Brivanlou, A.H., et al. (2010). APCDD1 is a novel Wnt inhibitor mutated in hereditary hypotrichosis simplex. Nature 464, 1043–1047. 10.1038/nature08875.

30. Zeng, S., Zhang, Y., Ma, J., Deng, G., Qu, Y., Guo, C., Han, Y., Yin, L., Cai, C., Li, Y., et al. (2017). BMP4 promotes metastasis of hepatocellular carcinoma by an induction of epithelial–mesenchymal transition via upregulating ID2. Cancer Lett. 390, 67–76. 10.1016/j.canlet.2016.12.042.

31. Aibar, S., González-Blas, C.B., Moerman, T., Huynh-Thu, V.A., Imrichova, H., Hulselmans, G., Rambow, F., Marine, J.-C., Geurts, P., Aerts, J., et al. (2017). SCENIC: Single-cell regulatory network inference and clustering. Nat Methods 14, 1083–1086. 10.1038/nmeth.4463.

32. Yang, L., Wang, W., Qiu, W., Guo, Z., Bi, E., and Xu, C. (2017). A single-cell transcriptomic analysis reveals precise pathways and regulatory mechanisms underlying hepatoblast differentiation. Hepatology 66, 1387–1401. 10.1002/hep.29353.

33. Kumari, S., Mukhopadhyay, G., and Tyagi, R.K. (2012). Transcriptional Regulation of Mouse PXR Gene: An Interplay of Transregulatory Factors. PLoS ONE 7, e44126. 10.1371/journal.pone.0044126.

34. Gougelet, A., Torre, C., Veber, P., Sartor, C., Bachelot, L., Denechaud, P., Godard, C., Moldes, M., Burnol, A., Dubuquoy, C., et al. (2014). T-cell factor 4 and β-catenin chromatin occupancies pattern zonal liver metabolism in mice. Hepatology 59, 2344–2357. 10.1002/hep.26924.

35. Behrens, J., Kries, J.P. von, Kühl, M., Bruhn, L., Wedlich, D., Grosschedl, R., and Birchmeier, W. (1996). Functional interaction of β-catenin with the transcription factor LEF-1. Nature 382, 638–642. 10.1038/382638a0.

36. Korinek, V., Barker, N., Morin, P.J., Wichen, D. van, Weger, R. de, Kinzler, K.W., Vogelstein, B., and Clevers, H. (1997). Constitutive Transcriptional Activation by a β-Catenin-Tcf Complex in APC−/− Colon Carcinoma. Science 275, 1784–1787. 10.1126/science.275.5307.1784.

37. Morin, P.J., Sparks, A.B., Korinek, V., Barker, N., Clevers, H., Vogelstein, B., and Kinzler, K.W. (1997). Activation of β-Catenin-Tcf Signaling in Colon Cancer by Mutations in β-Catenin or APC. Science 275, 1787–1790. 10.1126/science.275.5307.1787.

38. Pagella, P., Söderholm, S., Nordin, A., Zambanini, G., Ghezzi, V., Jauregi-Miguel, A., and Cantù, C. (2023). The time-resolved genomic impact of Wnt/β-catenin signaling. Cell Syst. 14, 563–581.e7. 10.1016/j.cels.2023.06.004.

39. Halpern, K.B., Shenhav, R., Matcovitch-Natan, O., Tóth, B., Lemze, D., Golan, M., Massasa, E.E., Baydatch, S., Landen, S., Moor, A.E., et al. (2017). Single-cell spatial reconstruction reveals global division of labour in the mammalian liver. Nature 542, 352–356. 10.1038/nature21065.

40. Aizarani, N., Saviano, A., Sagar, Mailly, L., Durand, S., Herman, J.S., Pessaux, P., Baumert, T.F., and Grün, D. (2019). A human liver cell atlas reveals heterogeneity and epithelial progenitors. Nature 572, 199–204. 10.1038/s41586-019-1373-2.

41. DeLaForest, A., Furio, F.D., Jing, R., Ludwig-Kubinski, A., Twaroski, K., Urick, A., Pulakanti, K., Rao, S., and Duncan, S.A. (2018). HNF4A Regulates the Formation of Hepatic Progenitor Cells from Human iPSC-Derived Endoderm by Facilitating Efficient Recruitment of RNA Pol II. Genes-basel 10, 21. 10.3390/genes10010021.

42. Park, W.S., Oh, R.R., Park, J.Y., Kim, P.J., Shin, M.S., Lee, J.H., Kim, H.S., Lee, S.H., Kim, S.Y., Park, Y.G., et al. (2001). Nuclear localization of β-catenin is an important prognostic factor in hepatoblastoma. J. Pathol. 193, 483–490. 10.1002/1096-9896(2000)9999:9999<::aid-path804>3.0.co;2-r.

43. Behrens, A., Sibilia, M., David, J., Möhle-Steinlein, U., Tronche, F., Schütz, G., and Wagner, E.F. (2002). Impaired postnatal hepatocyte proliferation and liver regeneration in mice lacking c-jun in the liver. Embo J 21, 1782–1790. 10.1093/emboj/21.7.1782.

44. Beck, A., Eberherr, C., Hagemann, M., Cairo, S., Häberle, B., Vokuhl, C., Schweinitz, D. von, and Kappler, R. (2016). Connectivity map identifies HDAC inhibition as a treatment option of high-risk hepatoblastoma. Cancer Biol Ther 17, 1168–1176. 10.1080/15384047.2016.1235664.

45. Kats, D., Ricker, C.A., Berlow, N.E., Noblet, B., Nicolle, D., Mevel, K., Branchereau, S., Judde, J.-G., Stiverson, C.D., Stiverson, C.L., et al. (2019). Volasertib preclinical activity in high-risk hepatoblastoma. Oncotarget 10, 6403–6417. 10.18632/oncotarget.27237.

46. Johnston, M.E., Rivas, M.P., Nicolle, D., Gorse, A., Gulati, R., Kumbaji, M., Weirauch, M.T., Bondoc, A., Cairo, S., Geller, J., et al. (2021). Olaparib Inhibits Tumor Growth of Hepatoblastoma in Patient-Derived Xenograft Models. Hepatology Baltim Md 74, 2201– 2215. 10.1002/hep.31919.

47. Bleijs, M., Pleijte, C., Engels, S., Ringnalda, F., Meyer-Wentrup, F., Wetering, M. van de, and Clevers, H. (2021). EWSR1-WT1 Target Genes and Therapeutic Options Identified in a Novel DSRCT In Vitro Model. Cancers 13, 6072. 10.3390/cancers13236072.

48. Meister, M.T., Koerkamp, M.J.A.G., Souza, T., Breunis, W.B., Frazer-Mendelewska, E., Brok, M., DeMartino, J., Manders, F., Calandrini, C., Kerstens, H.H.D., et al. (2022). Mesenchymal tumor organoid models recapitulate rhabdomyosarcoma subtypes. EMBO Mol. Med. 14, e16001. 10.15252/emmm.202216001.

49. López-Terrada, D., Alaggio, R., Dávila, M.T. de, Czauderna, P., Hiyama, E., Katzenstein, H., Leuschner, I., Malogolowkin, M., Meyers, R., Ranganathan, S., et al. (2014). Towards an international pediatric liver tumor consensus classification: proceedings of the Los Angeles COG liver tumors symposium. Modern Pathol 27, 472–491. 10.1038/modpathol.2013.80.

50. Gebhardt, R., and Hovhannisyan, A. (2010). Organ patterning in the adult stage: The role of Wnt/β-catenin signaling in liver zonation and beyond. Dev. Dyn. 239, 45–55. 10.1002/dvdy.22041.

51. Preziosi, M., Okabe, H., Poddar, M., Singh, S., and Monga, S.P. (2018). Endothelial Wnts regulate β-catenin signaling in murine liver zonation and regeneration: A sequel to the Wnt– Wnt situation. Hepatol. Commun. 2, 845–860. 10.1002/hep4.1196.

52. Annunziato, S., Sun, T., and Tchorz, J.S. (2022). The RSPO-LGR4/5-ZNRF3/RNF43 module in liver homeostasis, regeneration, and disease. Hepatology 76, 888–899. 10.1002/hep.32328.

53. Watt, A.J., Garrison, W.D., and Duncan, S.A. (2003). HNF4: A central regulator of hepatocyte differentiation and function. Hepatology 37, 1249–1253. 10.1053/jhep.2003.50273.

54. DeLaForest, A., Nagaoka, M., Si-Tayeb, K., Noto, F.K., Konopka, G., Battle, M.A., and Duncan, S.A. (2011). HNF4A is essential for specification of hepatic progenitors from human pluripotent stem cells. Development 138, 4143–4153. 10.1242/dev.062547.

55. Parviz, F., Matullo, C., Garrison, W.D., Savatski, L., Adamson, J.W., Ning, G., Kaestner, K.H., Rossi, J.M., Zaret, K.S., and Duncan, S.A. (2003). Hepatocyte nuclear factor 4α controls the development of a hepatic epithelium and liver morphogenesis. Nat. Genet. 34, 292–296. 10.1038/ng1175.

56. DeLaForest, A., Nagaoka, M., Si-Tayeb, K., Noto, F.K., Konopka, G., Battle, M.A., and Duncan, S.A. (2011). HNF4A is essential for specification of hepatic progenitors from human pluripotent stem cells. Development 138, 4143–4153. 10.1242/dev.062547.

57. Gougelet, A., Torre, C., Veber, P., Sartor, C., Bachelot, L., Denechaud, P., Godard, C., Moldes, M., Burnol, A., Dubuquoy, C., et al. (2014). T-cell factor 4 and β-catenin chromatin occupancies pattern zonal liver metabolism in mice. Hepatology 59, 2344–2357. 10.1002/hep.26924.

58. Yang, M., Li, S.-N., Anjum, K.M., Gui, L.-X., Zhu, S.-S., Liu, J., Chen, J.-K., Liu, Q.-F., Ye, G.-D., Wang, W.-J., et al. (2013). A double-negative feedback loop between Wnt–β-catenin signaling and HNF4α regulates epithelial–mesenchymal transition in hepatocellular carcinoma. J. Cell Sci. 126, 5692–5703. 10.1242/jcs.135053.

59. Billin, A.N., Thirlwell, H., and Ayer, D.E. (2000). β-Catenin–Histone Deacetylase Interactions Regulate the Transition of LEF1 from a Transcriptional Repressor to an Activator. Mol Cell Biol 20, 6882–6890. 10.1128/mcb.20.18.6882-6890.2000.

60. Chi, A., Remick, S., and Tse, W. (2013). EGFR inhibition in non-small cell lung cancer: current evidence and future directions. Biomark Res 1, 2. 10.1186/2050-7771-1-2.

61. Kommalapati, A., Tella, S.H., Borad, M., Javle, M., and Mahipal, A. (2021). FGFR Inhibitors in Oncology: Insight on the Management of Toxicities in Clinical Practice. Cancers 13, 2968. 10.3390/cancers13122968.

62. Suraweera, A., O’Byrne, K.J., and Richard, D.J. (2018). Combination Therapy With Histone Deacetylase Inhibitors (HDACi) for the Treatment of Cancer: Achieving the Full Therapeutic Potential of HDACi. Frontiers Oncol 8, 92. 10.3389/fonc.2018.00092.

63. Maouche, N., Kishore, B., Bhatti, Z., Basu, S., Karim, F., Sundararaman, S., Collings, F., Tseu, B., Leary, H., Ryman, N., et al. (2022). Panobinostat in combination with bortezomib and dexamethasone in multiply relapsed and refractory myeloma; UK routine care cohort. PLoS ONE 17, e0270854. 10.1371/journal.pone.0270854.

64. Wagner, A., Schwarzmayr, T., Häberle, B., Vokuhl, C., Schmid, I., Kaller, M., Hermeking, H., Schweinitz, D. von, and Kappler, R. (2020). SP8 Promotes an Aggressive Phenotype in Hepatoblastoma via FGF8 Activation. Cancers 12, 2294. 10.3390/cancers12082294.

65. Saltsman, J.A., Hammond, W.J., Narayan, N.J.C., Requena, D., Gehart, H., Lalazar, G., LaQuaglia, M.P., Clevers, H., and Simon, S. (2020). A Human Organoid Model of Aggressive Hepatoblastoma for Disease Modeling and Drug Testing. Cancers 12, 2668. 10.3390/cancers12092668.

66. Wu, P.V., and Nusse, R. (2022). Hepatocytes, Methods and Protocols. Methods Mol Biology 2544, 259–267. 10.1007/978-1-0716-2557-6_19.

67. Peng, W.C., Logan, C.Y., Fish, M., Anbarchian, T., Aguisanda, F., Álvarez-Varela, A., Wu, P., Jin, Y., Zhu, J., Li, B., et al. (2018). Inflammatory Cytokine TNFα Promotes the Long-Term Expansion of Primary Hepatocytes in 3D Culture. Cell 175, 1607–1619.e15. 10.1016/j.cell.2018.11.012.

68. Miao, Y., Ha, A., Lau, W. de, Yuki, K., Santos, A.J.M., You, C., Geurts, M.H., Puschhof, J., Pleguezuelos-Manzano, C., Peng, W.C., et al. (2020). Next-Generation Surrogate Wnts Support Organoid Growth and Deconvolute Frizzled Pleiotropy In Vivo. Cell Stem Cell. 10.1016/j.stem.2020.07.020.

69. Kluiver, T.A., Kraaier, L.J., and Peng, W.C. (2022). Hepatocytes, Methods and Protocols. Methods Mol Biology 2544, 1–13. 10.1007/978-1-0716-2557-6_1.

70. Hao, Y., Hao, S., Andersen-Nissen, E., Mauck, W.M., Zheng, S., Butler, A., Lee, M.J., Wilk, A.J., Darby, C., Zager, M., et al. (2021). Integrated analysis of multimodal single-cell data. Cell 184, 3573–3587.e29. 10.1016/j.cell.2021.04.048.

71. Yang, S., Corbett, S.E., Koga, Y., Wang, Z., Johnson, W.E., Yajima, M., and Campbell, J.D. (2020). Decontamination of ambient RNA in single-cell RNA-seq with DecontX. Genome Biol 21, 57. 10.1186/s13059-020-1950-6.

72. Calandrini, C., Schutgens, F., Oka, R., Margaritis, T., Candelli, T., Mathijsen, L., Ammerlaan, C., Ineveld, R.L. van, Derakhshan, S., Haan, S. de, et al. (2020). An organoid biobank for childhood kidney cancers that captures disease and tissue heterogeneity. Nat Commun 11, 1310. 10.1038/s41467-020-15155-6.

73. Slyper, M., Porter, C.B.M., Ashenberg, O., Waldman, J., Drokhlyansky, E., Wakiro, I., Smillie, C., Smith-Rosario, G., Wu, J., Dionne, D., et al. (2020). A single-cell and single-nucleus RNA-Seq toolbox for fresh and frozen human tumors. Nat Med 26, 792–802. 10.1038/s41591-020-0844-1.

74. Butler, A., Hoffman, P., Smibert, P., Papalexi, E., and Satija, R. (2018). Integrating single-cell transcriptomic data across different conditions, technologies, and species. Nat Biotechnol 36, 411–420. 10.1038/nbt.4096.

75. Stuart, T., Butler, A., Hoffman, P., Hafemeister, C., Papalexi, E., Mauck, W.M., Hao, Y., Stoeckius, M., Smibert, P., and Satija, R. (2019). Comprehensive Integration of Single-Cell Data. Cell 177, 1888–1902.e21. 10.1016/j.cell.2019.05.031.

