## Supplemental file for "Multi-dimensional profiling of hepatoblastomas and patient-derived tumor organoids uncovers tumor subpopulations with divergent WNT activation profiles and identifies pan-hepatoblastoma drug sensitivities"

**Supplemental information**

**Table S1. Overview of patients, tissues and organoid models described in this study.**

| | | | | | | | | **Tissue** | | | **Organoids** | | | | |
|---|---|---|---|---|---|---|---|---|---|---|---|---|---|---|---|
| **Patient** | **Tumor** | **Risk** | **Age** | **Sex** | **PRETEXT** | ***CTNNB1* (tumor AF%)** | **Material** | **Spatial** | **IF** | **scRNA** | **Organoid** | **scRNA** | **Drug screen** | **Medium** | **Sanger *CTNNB1*** |
| PT2 | HB | High | 2-8y | M | II | Exon 3 del | Post-chemo | Yes | | | | | | | |
| PT3 | HB | High | <2y | F | II | p.T41A (36%) | Pre-chemo | | | | 3E | Yes | Yes | Full | Matched |
| PT8 | HB | Low | <2y | F | II | Exon 3 del | Pre-chemo | | | | 8F | Yes | Yes | Full | |
| PT9 | HB | High | <2y | M | III | p.G34V (39%) | Pre-chemo | | Yes | | | | | | |
| | | | | | | | Post-chemo | | Yes | | | | | | |
| PT10 | HB | Intermediate | <2y | F | III | Exon 3 del | Pre-chemo | | Yes | | 10F | Yes | Yes | Full | Matched |
| PT13 | HB | Very low | <2y | M | II | p.D32N (40%) | Pre-chemo | | | | 13F | Yes | Yes | Full | Matched |
| | | | | | | | Pre-chemo | Yes | Yes | Yes | 13E | Yes | Yes | Full | |
| PT14 | HB | Intermediate | 2-8y | F | IV | p.S23_G34del (31%) | Pre-chemo | | Yes | | | | | | |
| | | | | | | | Post-chemo | Yes | | | | | | | |
| PT15 | HB | High | 2-8y | M | IV | p.S29F; p.D32Y (37%) | Pre-chemo | | Yes | | | | | | |
| PT16 | HB | Intermediate | <2y | M | II | Exon 3 del | Post-chemo | Yes | | | | | | | |
| PT17 | HB | High | <2y | F | II | p.V22_Q78delinsE | Pre-chemo | | Yes | | 17E | Yes | Yes | Full | Matched |
| | | | | | | | Post-chemo | | Yes | | 17F | Yes | Yes | Reduced | Matched |
| PT20 | HB | High | 2-8y | M | IV | Exon 3 del | Pre-chemo | | Yes | | | | | | |
| PT22 | HB | Intermediate | <2y | F | I | Exon 3 del | Pre-chemo | | Yes | | 22E | Yes | Yes | Full | Matched |
| PT27 | HB | High | 2-8y | M | IV | Exon 3 del | Pre-chemo | | | | 27 | No | No | Reduced | |
| PT96 | HB | UNK | UNK | UNK | UNK | UNK | Post-chemo | | | | 96F | Yes | Yes | Reduced | Exon 3 del |
| PT121 | HB | UNK | UNK | UNK | UNK | UNK | Relapse (lung met) | | | | 121E | Yes | Yes | Full | p.D32N |
| PT135 | HB | UNK | UNK | UNK | UNK | UNK | Relapse | | | | 135 | Yes | Yes | Full | p.S29F; p.D32Y |

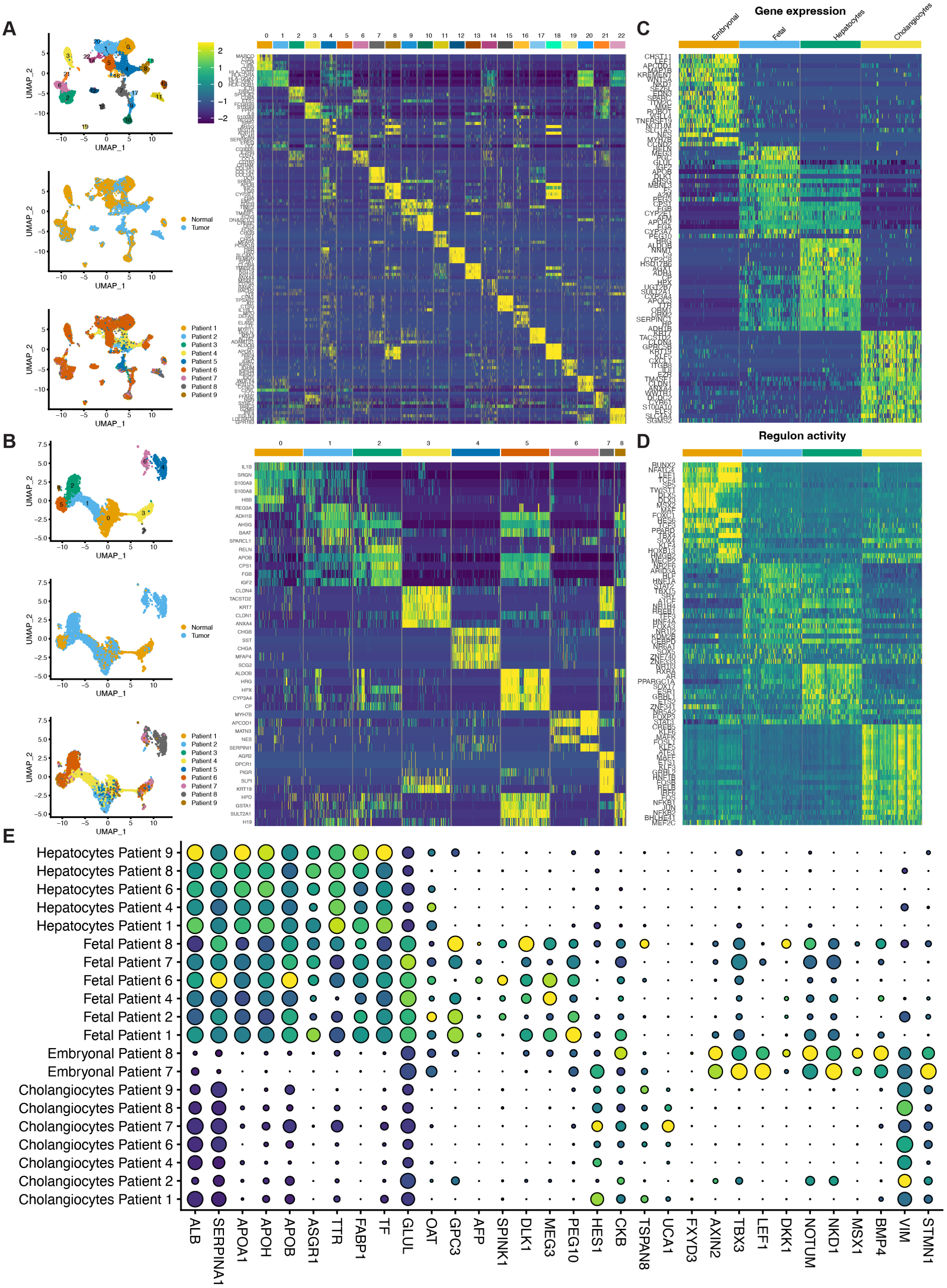


**Figure S1. Single cell hepatoblastoma processing and analysis.**
(A) UMAP showing the full dataset from *Song et al.* after data filtering, normalization, scaling, DecontX, fastMNN batch correction, dimensional reduction, and clustering (left). Heatmap showing the top 5 differentially expressed markers per cluster (right). The epithelial normal and tumor clusters 4, 8, 11, 13 and 18 were subsetted and unbiased clustering was performed.

(B) UMAP showing the reprocessed subsetted clusters (left). Heatmap showing the top 5 differentially expressed markers per cluster of the subsetted object (right). We removed low-quality clusters likely contaminated with non-parenchymal cells, and the neuroendocrine cluster (4) for the final object. Clusters 2, 3, 5, 6, 7 and 8 were retained.
(C) Heatmap showing top differentially expressed genes of the final clusters.
(D) Single-cell heatmap showing differentially active transcription factor regulons, using SCENIC, for each of the tumor clusters as well as normal hepatocytes and cholangiocytes.
(E) Dot plot showing selected marker expression per cluster, per patient. Only populations with at least 10 cells are shown.

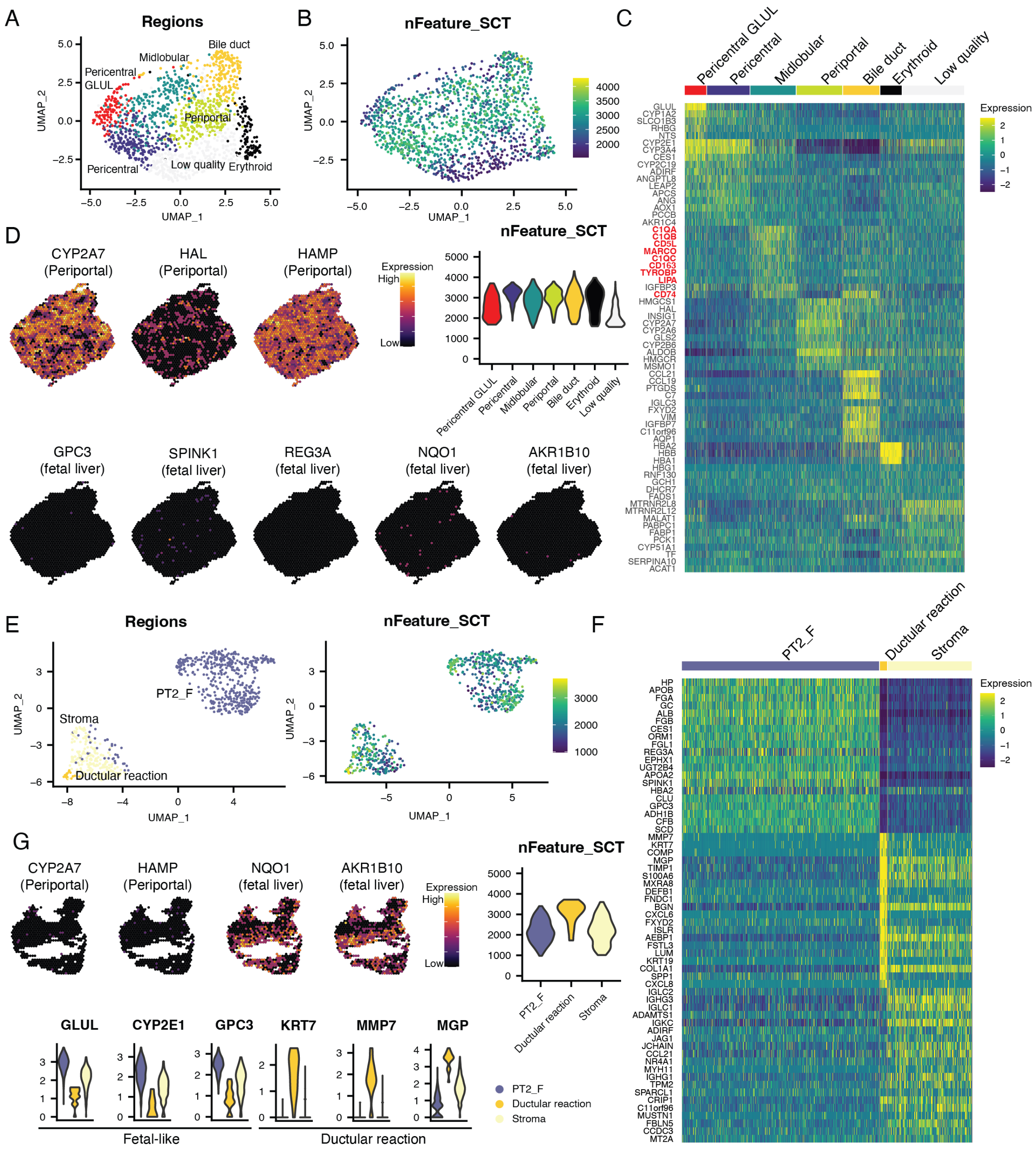

A
Regions
Pericentral GLUL
Midlobular
Bile duct
Periportal
Pericentral
Low quality
Erythroid
UMAP_1
UMAP_2
B
nFeature_SCT
C
Pericentral GLUL
Pericentral
Midlobular
Periportal
Bile duct
Erythroid
Low quality
Expression
D
CYP2A7 (Periportal)
HAL (Periportal)
HAMP (Periportal)
Expression
High
Low
nFeature_SCT
GPC3 (fetal liver)
SPINK1 (fetal liver)
REG3A (fetal liver)
NQO1 (fetal liver)
AKR1B10 (fetal liver)
E
Regions
PT2_F
Stroma
Ductular reaction
nFeature_SCT
F
PT2_F
Ductular reaction
Stroma
Expression
G
CYP2A7 (Periportal)
HAMP (Periportal)
NQO1 (fetal liver)
AKR1B10 (fetal liver)
Expression
High
Low
nFeature_SCT
GLUL
CYP2E1
GPC3
KRT7
MMP7
MGP
Fetal-like
Ductular reaction
PT2_F
Ductular reaction
Stroma

H

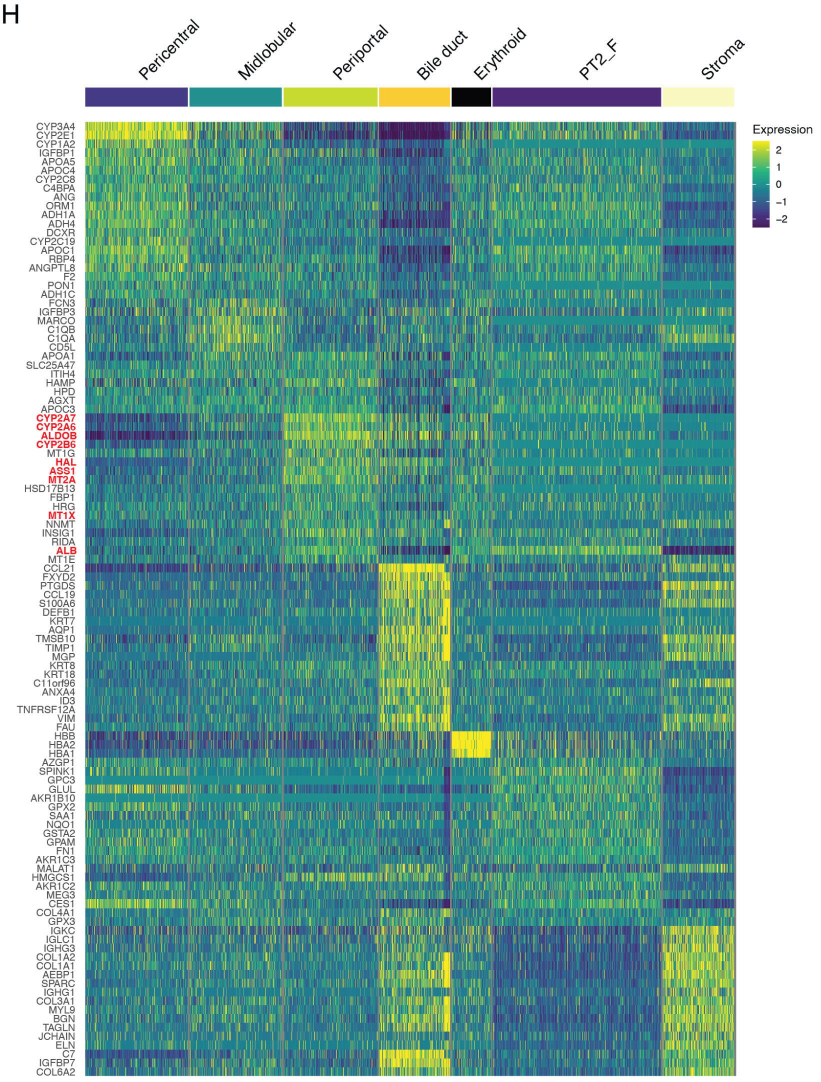


**Figure S2. Spatial transcriptomic analysis of PT2 normal liver and fetal hepatoblastoma.**

(A) Unsupervised graph-based clustering was performed on the ST spots of the distal normal liver tissue from PT2, identifying *e.g,* GLUL pericentral, pericentral, midlobular, periportal and bile duct regions.

(B) Quality control was performed based on number of SCTransformed features per cluster/spot (“nFeatures_SCT”>2000), visualized in UMAP representation and violin plot. One cluster was assigned as “Low quality” and was excluded form downstream analyses.

(C) Heatmap of differentially expressed genes between the liver zones. Of note, the midlobular region showed high expression of genes associated with macrophages (marked in red).

(D) Spatial plots showing expression of an additional periportal marker and absence of fetal liver markers in distal normal liver tissue.

(E) Violin plot and UMAP representation of number of SCTransformed features per cluster/spot. In the stroma region the cell density is likely lower based on lower number of features per spot.

(F) Heatmap of differentially expressed genes between the clusters. Tumor spots expressed pericentral hepatocyte and fetal liver markers. Stromal regions expressed immune and endothelial markers (*CCL21*, *CCL19, MGP*).

(G) Spatial plots showing absence of an additional periportal marker and expression of fetal liver markers in tumor regions. Violin plots of fetal hepatoblastoma markers and cholangiocyte and portal area markers, identifying ductular reaction in the tumor stroma region.

(H) Heatmap of differentially expressed genes between the clusters of the PT2 tumor and distal normal sections. Tumor spots expressed pericentral hepatocyte and fetal liver markers but showed reduced expression of periportal markers (marked in red).

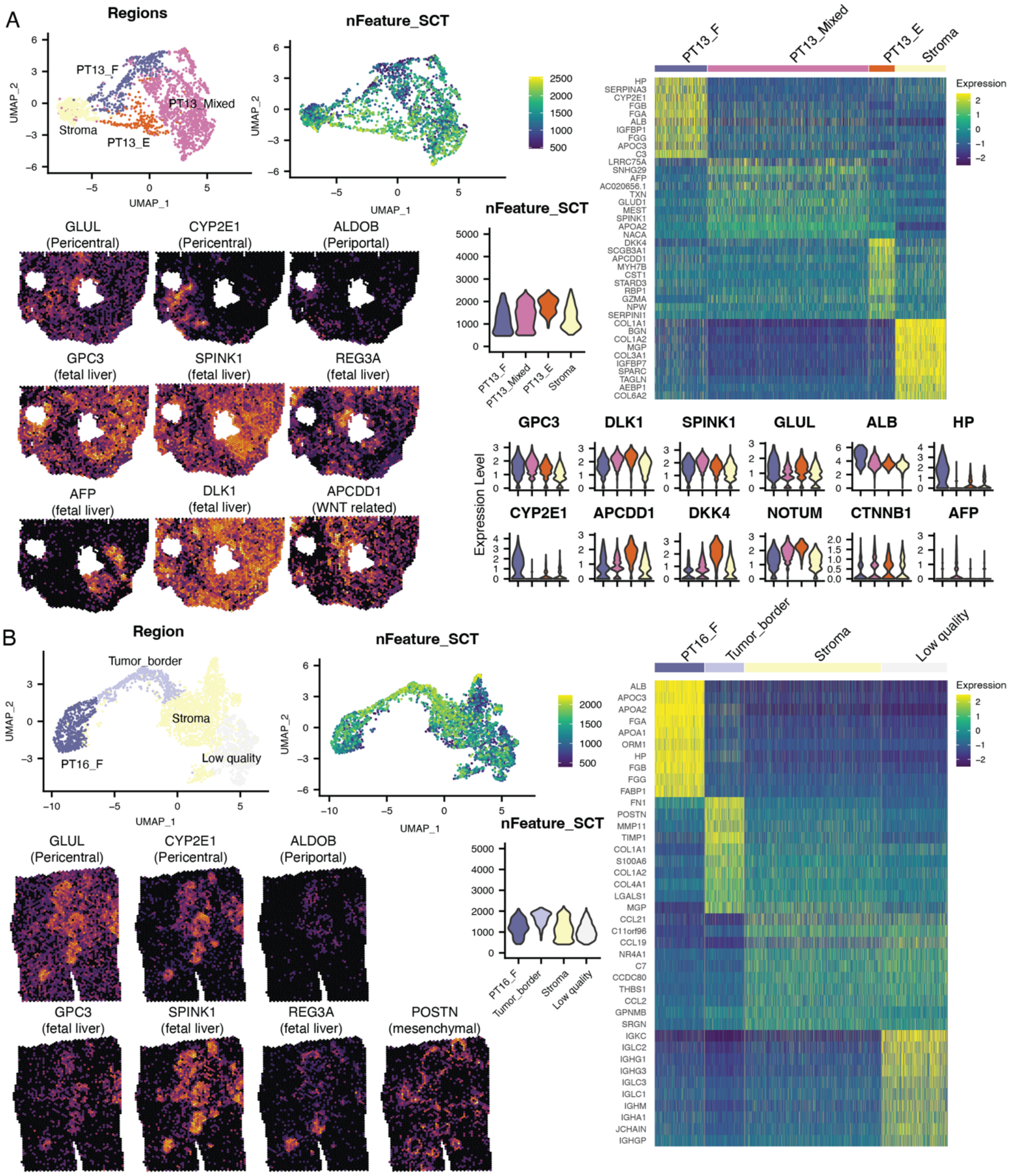
A
Regions
PT13_F
PT13_Mixed
Stroma
PT13_E
UMAP_1
UMAP_2
nFeature_SCT
2500
2000
1500
1000
500
Expression
GLUL
(Pericentral)
CYP2E1
(Pericentral)
ALDOB
(Periportal)
GPC3
(fetal liver)
SPINK1
(fetal liver)
REG3A
(fetal liver)
AFP
(fetal liver)
DLK1
(fetal liver)
APCDD1
(WNT related)
HP
SERPINA3
CYP2E1
FGB
FGA
ALB
IGFBP1
FGG
APOC3
C3
LRRC75A
SNHG29
AFP
AC020656.1
TXN
GLUD1
MEST
SPINK1
APOA2
NACA
DKK4
SCGB3A1
APCDD1
MYH7B
CST1
STARD3
RBP1
GZMA
NPW
SERPINI1
COL1A1
BGN
COL1A2
MGP
COL3A1
IGFBP7
SPARC
TAGLN
AEBP1
COL6A2
Expression Level
GPC3
DLK1
SPINK1
GLUL
ALB
HP
CYP2E1
APCDD1
DKK4
NOTUM
CTNNB1
AFP
B
Region
Tumor_border
Stroma
PT16_F
Low quality
nFeature_SCT
GLUL
(Pericentral)
CYP2E1
(Pericentral)
ALDOB
(Periportal)
GPC3
(fetal liver)
SPINK1
(fetal liver)
REG3A
(fetal liver)
POSTN
(mesenchymal)
ALB
APOC3
APOA2
FGA
APOA1
ORM1
HP
FGB
FGG
FABP1
FN1
POSTN
MMP11
TIMP1
COL1A1
S100A6
COL1A2
COL4A1
LGALS1
MGP
CCL21
C11orf96
CCL19
NR4A1
C7
CCDC80
THBS1
CCL2
GPNMB
SRGN
IGKC
IGLC2
IGHG1
IGHG3
IGLC3
IGLC1
IGHM
IGHA1
JCHAIN
IGHGP

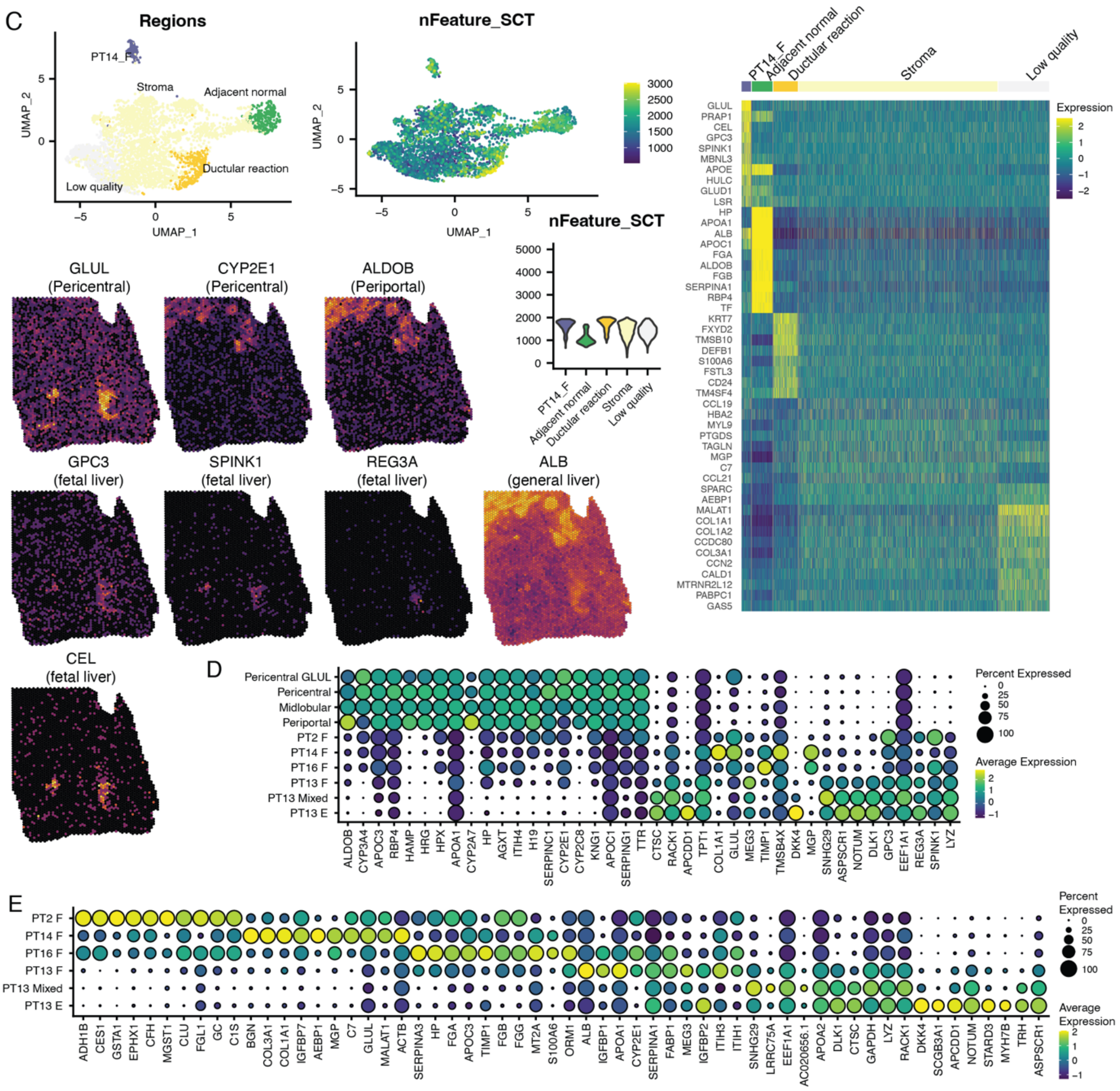


**Figure S3. Spatial transcriptomic analysis of three additional hepatoblastoma tissues.**

For each section individually, we inspected the H&E stainings and performed: (1) unsupervised graph-based clustering, (2) quality control assessment (SCTransformed features per cluster/spot visualized in UMAP representation and violin plot), (3) differential gene expression analysis (heatmap visualizing the top 10 differentially expressed genes per cluster) and (4) marker gene expression (visualized in violin plots or spatial distribution).

(A) The tumor of PT13 was resected prior to chemotherapy. High expression of fetal liver markers could be observed throughout the tumor regions. At least three different tumor clusters were identified; fetal-like (higher expression of hepatocyte genes), embryonal-like (higher expression of WNT related genes), and regions likely containing a mix of both populations of tumor cells. Additional tissue heterogeneity could be observed based on markers such as *AFP*.

(B) The tumor of PT16 contained fetal-like tumor regions, with expression of pericentral hepatocyte and fetal liver markers. The tumor border showed a distinct expression profile, with high levels of *POSTN* (mesenchymal marker) and *KRT19* (cholangiocyte marker).

(C) The tumor of PT14 contained fetal-like tumor, normal liver, and stroma. Adjacent normal liver showed higher expression of hepatocyte markers compared to the fetal-like tumor regions, which had higher expression of fetal liver markers. One of the stromal regions showed expression of *KRT7* (cholangiocyte marker), therefore likely containing both stromal cells and ductal cells. On the H&E staining, ductular reaction could be confirmed, which is frequently observed at the border between normal liver and hepatoblastoma.

(D) Dot plot of the top 20 differentially expressed genes between distal hepatocytes and the combined tumor regions. There is an increased expression of fetal liver genes and a decreased expression of periportal markers in the tumor regions. Inter-tumor heterogeneity can also be observed.

(E) Dot plot of the top 10 differentially expressed genes between the tumor clusters, illustrating tumor-specific expression profiles, and additional heterogeneity within the PT13 tumor.

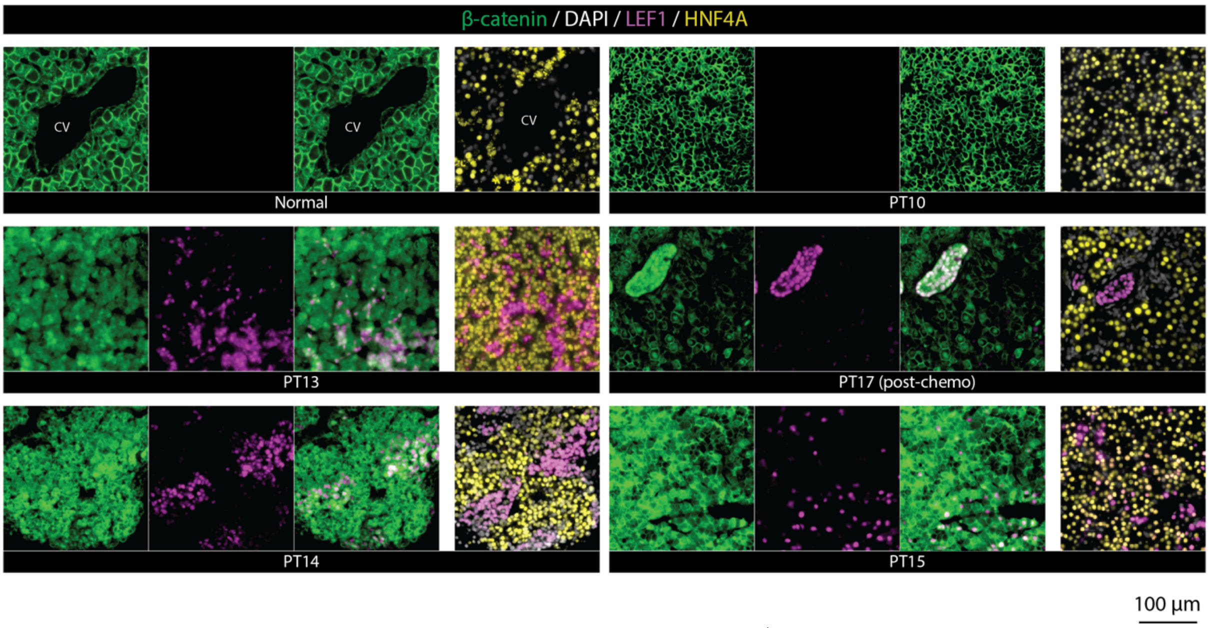


**Figure S4. Β-catenin stains more nuclear rather than membranous in LEF+ hepatoblastoma cells.**
Co-staining of β-catenin and LEF in normal liver and tumors with fetal and embryonal regions, as well as co-staining of HNF4A and LEFF1 in serial sections are shown. Β-catenin demonstrated heterogeneity in its staining pattern compared to LEF1.

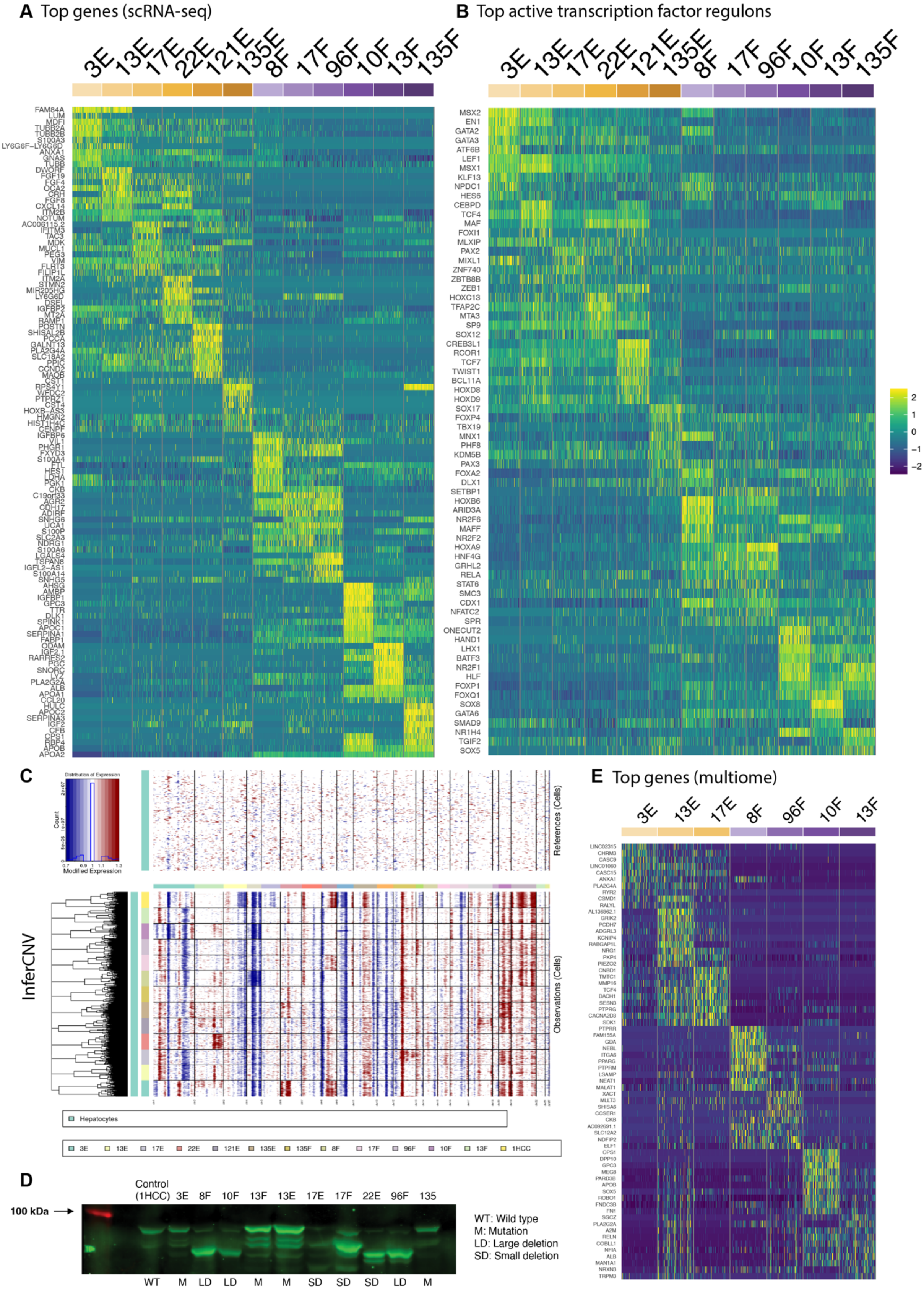


**Figure S5. Single cell HB organoid analysis.**
(A) Heatmap showing the top differentially expressed genes per organoid model.
(B) Heatmap showing the top active gene regulatory networks per organoid model using SCENIC reconstructed regulons.
(C) InferCNV based on organoid scRNA-seq data confirms the chromosomal copy number variations of the primary tumors are maintained in the organoids. The hepatocyte cluster from the Song *et al.* dataset was used as reference cells.

(D) Western blot confirms expression of mutant β-catenin proteins in organoid samples with exon 3 deletions. As positive control, organoid samples with missense mutations and an HCC organoid sample with wild type *CTNNB1* were used.
(E) Heatmap showing the top differentially expressed genes per organoid model based on the RNA counts of the single nucleus multiome dataset.

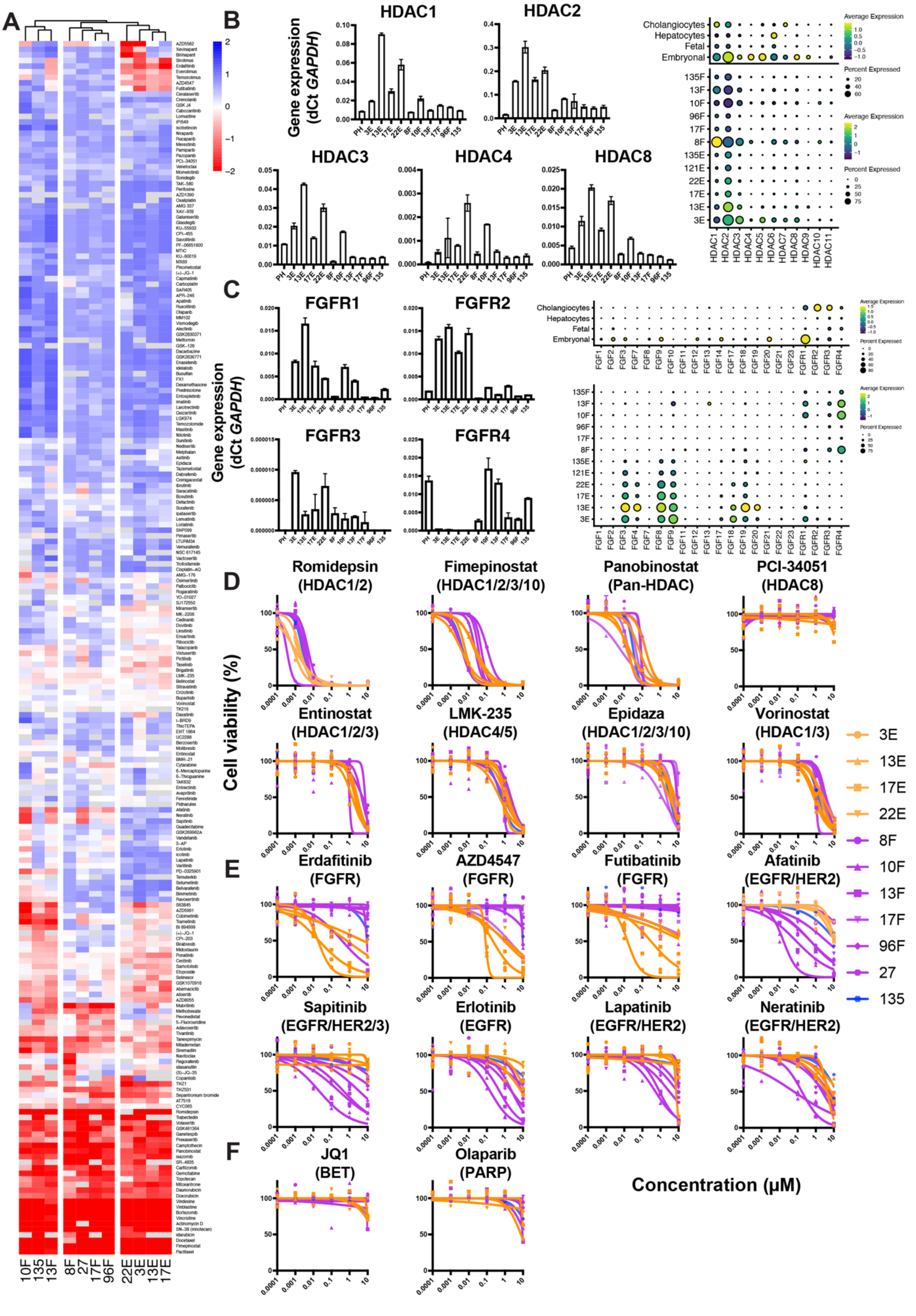


**Figure S6. High-throughput drug screening of hepatoblastoma organoids.**
(A) Full, scaled heatmap showing AUC values for dose response curves for all compounds tested.

(B) qRT-PCR graphs showing expression of HDAC genes in different organoid models normalized against *GAPDH* (left). Error bars respresent standard deviations. Dot plots showing expression of all HDAC genes in organoids and tissues as measured by scRNA-seq (right).
(C) qRT-PCR graphs showing expression of FGFR genes in different organoid models normalized against *GAPDH* (left). Error bars respresent standard deviations. Dot plots showing expression of all HDAC genes in organoids and tissues as measured by scRNA-seq (right).
(D) Drug screening dose response curves for selected HDAC inhibitors. Points represent two technical replicates.
(E) Drug screening dose response curves for selected FGFR and EGFR inhibitors. Points represent two technical replicates.
(F) Drug screening dose response curves for JQ1 and Olaparib, which have little effect in our screen.

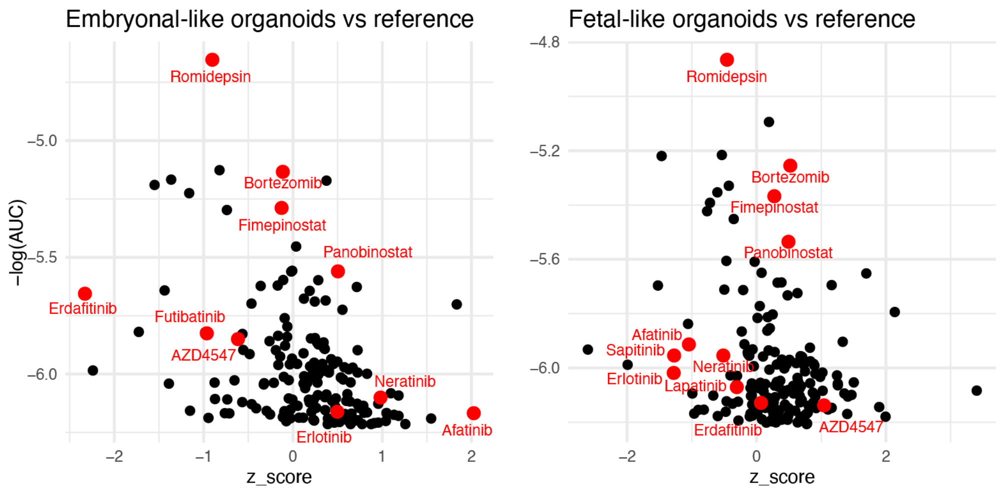


**Figure S7. Hepatoblastoma organoid drug responses compared to the pediatric tumor reference cohort.** Volcano plots showing the z-scores of fetal-like (left) and embryonal-like (right) drug responses (AUC values) versus the pediatric tumor organoid reference cohort. Lower z-scores indicate more specific sensitivity for hepatoblastoma organoids. On the y-axis, the negative logarithm of the average AUC value for the respective hepatoblastoma organoids is plotted. Only drugs with IC50 values are shown.
